# Segmental copy number amplifications are more stable than aneuploidies in the absence of selection

**DOI:** 10.1101/2025.07.21.665951

**Authors:** Titir De, Nadav Ben Nun, Pieter Spealman, Ina Suresh, Grace Avecilla, Farah Abdul-Rahman, Yoav Ram, David Gresham

## Abstract

Copy number variants (CNVs) are DNA duplications and deletions that cause genetic variation, underlying rapid adaptive evolution. CNVs often confer selective advantages, but can also incur fitness costs. Evolution of *Saccharomyces cerevisiae* in nutrient-limited chemostats recurrently selects for amplifications of nutrient transporter genes. However, their fate upon return to a non-selective environment remains unknown. To investigate CNV fitness and stability upon removing the original selection pressure, we studied 15 CNV lineages (11 segmental, 4 whole-chromosomal amplifications) selected in nitrogen-limited chemostats. CNV stability was monitored using fluorescent reporters during propagation in nutrient-rich batch cultures for 110–220 generations. All aneuploid lineages showed rapid CNV loss and reversion to a single-copy genotype, whereas segmental amplifications were remarkably stable– one of 11 strains reverted. Pairwise fitness competitions in rich media revealed strong fitness defects associated solely with CNVs that reverted; reversion led to increased fitness. Using simulation-based inference to estimate reversion rates and fitness effects, we determined negative selection as the primary driver of CNV loss. Whole-genome sequencing revealed that reversion of aneuploids and a segmental amplification left no evidence of prior CNV existence, rendering revertant genomes indistinguishable from the single-copy ancestor. Detailed characterization of a partial revertant identified chromosomal translocation, suggesting that extant CNVs can undergo structural diversification. Our findings provide novel evidence that most segmental CNVs adapted to nitrogen limitation are stable upon removal of selection, but costly gene amplifications are readily reversible. Together, these highlight the importance of CNVs in both long-term genome evolution and rapid, reversible adaptation to transient selection.

## Introduction

Understanding the determinants and dynamics of adaptation is a fundamental goal of evolutionary biology. Genetic differences within a species, acted upon by forces of natural selection, are a major driver of adaptive evolution. One prevalent source of natural genetic variation within species are copy number variants (CNVs) – duplications or deletions of DNA sequences [Iskow et al. 2012, Clop et al. 2012, Żmieńko et al. 2014, Zarrei et al. 2015]. CNVs are a distinct class of genetic variation which, unlike single nucleotide variations, typically involve large genomic regions, ranging from a few base pairs (bp) to an entire chromosome, and thus can affect multiple genes [Connolly et al. 2014]. Environmental conditions can lead to selection pressures that act upon CNVs within a population, selecting for those that are best adapted to that condition.

CNVs cause rapid phenotypic diversification by changing the amount of DNA template, thus modifying mRNA and protein levels. This can confer benefits that help the organism survive in a particular environment– such CNVs are rapidly selected for and often attain fixation in the population [Kondrashov 2012, Myhre et al. 2013, Robinson et al. 2023]. For example, CNVs have been found to underlie drug resistance and enhanced nutrient transport in microbes, as well as defence mechanisms against diseases in plants [Elde et al. 2012, Greenblum et al. 2015, Iantorno et al. 2017, Dolatabadian et al. 2017, Cowell et al. 2018]. CNVs can also aid in the survival and proliferation of cancer cells, through amplification of oncogenes, heightened adaptability, and faster resistance to anti-cancer treatments [Rutledge et al. 2016, Ben-David and Amon 2020, Lukow et al. 2021]. In addition to short-term effects, CNVs also mediate long-term evolution through neofunctionalisation and subfunctionalisation of functionally redundant gene copies [Ohno 1970, Innan and Kondrashov 2010, Freeling et al. 2015]. On the other hand, CNVs can also confer negative impacts e.g., duplications or deletions of key genes are involved in a wide range of human disorders such as haemophilia, Alzheimer’s disease, autism and schizophrenia. [Zhang et al. 2009]. Notably, the same CNV can potentially be beneficial, neutral, or deleterious depending on the relationship between its functional effect and the environment. Although there have been multiple studies of the dynamics, mechanisms, and effects of CNVs in selective conditions, the stability and maintenance of CNVs in populations once the selective pressure has been removed remains poorly understood.

CNVs are often associated with both fitness costs and fitness benefits [Tang and Amon 2013, Gerstein and Berman 2015]. CNVs are advantageous, and subject to positive selection, only when the benefits outweigh the costs in a particular environment. Typically, CNVs follow the ‘driver-hitchhiker’ model [Smith and Haigh 1974, Golzio and Katsanis 2013], wherein one of the amplified genes (driver) confers a fitness benefit, but other genes included in the amplified genomic region (hitchhikers) are neutral or confer fitness costs. These costs can be due to different reasons. First, the dosage burden model posits that the fitness cost is proportional to the elevation in unnecessary gene expression resulting from the CNV [Dekel and Alon 2005, Makanae et al. 2013, Bonney et al. 2015]. The fitness costs of unnecessary gene expression is evident in several studies, e.g., sterility is positively selected for in yeast during asexual growth due to elimination of unnecessary gene expression [Payen et al. 2016, Rice & McLysaght 2017] and experimental overexpression of individual genes shows clear evidence for deleterious effects [Robinson et al. 2021]. Second, the dosage imbalance model posits that fitness costs are derived from stoichiometric imbalances of dosage-sensitive genes [Wagner 2005, Veitia et al. 2008, Rice and McLysaght 2017]. Third, there may also be fitness costs due to stress associated with the higher levels of DNA replication, transcription, and translation that result from CNVs. Although prior research has shed light on the fitness benefits associated with the emergence and selection of CNVs, we do not know how the trade-offs between fitness costs and benefits influence their stability in the absence of the original selection conditions under which they arose.

Long-term experimental evolution allows the study of the evolutionary dynamics under controlled, replicated selection conditions [Lenski et al. 1991, Good et al. 2017]. An integral component of cell growth regulation is acquisition of essential nutrients from the environment through specialized nutrient transporters. When cells are grown continuously in a nutrient-limited environment in a chemostat [Gresham and Dunham 2014, Gresham and Hong 2015], *de novo* CNVs containing nutrient transporter genes are positively selected, as increased abundance of transporter proteins likely allows increased uptake of the limited nutrient [Spealman et al., 2025]. This evolutionary outcome has been shown in multiple studies and species: *Saccharomyces cerevisiae* under limitation of glucose, phosphate, sulfur, and nitrogen [Brown et al. 1998, Dunham et al. 2002, Gresham et al. 2008, Kao and Sherlock 2008, Hong and Gresham 2014, Payen et al. 2016], *Escherichia coli* limited for lactose [Horiuchi et al., 1963, Zhong et al. 2004], and *Salmonella typhimurium* in different carbon source limitations [Sonti and Roth, 1989]. Previous studies have shown that evolution in glutamine-limited chemostats repeatedly selects for *de novo* amplifications of *GAP1*, a general amino acid permease [Lauer et al. 2018, Avecilla et al. 2023, Chuong et al. 2025]. *GAP1* CNVs are also found in natural populations, such as in the nectar yeast *Metschnikowia reukaufii*, where multiple copies of *GAP1* are beneficial when amino acids are limited [Dhami et al. 2016]. Selection for CNVs containing nitrogen transporters is substrate specific. *MEP2* is a high-affinity ammonium transporter that is amplified under ammonium limitation, whereas *PUT4* is a proline transporter amplified by proline-limited selection [Gresham et al. 2010, Hong & Gresham 2014, Hong et al. 2018]. Although multiple studies have demonstrated the recurrent emergence and fixation of CNVs in nutrient-limited chemostats, it is unknown whether acquired CNVs are maintained or lost when the organism is returned to nutrient-replete conditions.

Quantifying the frequency of CNVs in heterogeneous populations using molecular methods poses a considerable challenge and is limited in throughput and resolution. Previously, we developed a method to visualise CNVs in evolving populations using a constitutively expressed fluorescent reporter gene inserted adjacent to the gene of interest [Lauer et al. 2018]. Since CNVs span large genomic regions, the gene of interest and the reporter are typically amplified or deleted together, allowing estimation of the frequency of CNVs on the basis of fluorescence levels [Lauer et al. 2018, Spealman et al. 2023].

In this study, we used budding yeast lineages that previously acquired nutrient transporter-associated genomic amplifications in chemostat evolution experiments, as a system to study the fitness costs and stability of CNVs in absence of the original selection pressure. Using a combination of long-term experimental evolution using serial batch culture in nutrient-rich media, and fitness assays of CNV and revertant lineages, we find that a majority of CNVs adapted to nitrogen limitation do not lead to significant fitness defects compared to the ancestral strain in absence of this selection pressure. As a result, most segmental CNVs are stable over hundreds of generations, despite not conferring a fitness benefit in the absence of the original selection pressure. By contrast, aneuploidies exhibit significant fitness costs and instability and therefore are rapidly lost in the absence of selection. The dynamics of aneuploidy loss differs between chromosomes; we find that chromosome XIV aneuploidy is lost much faster and ubiquitously whereas loss of chromosome XI aneuploidy is much slower and variable between populations. This difference, despite having the same copy number and similar chromosome sizes, is consistent with fitness costs of aneuploids depending on the specific hitchhiker genes on the chromosome. Although it appears to be rare for CNVs adapted to different kinds of nitrogen limitation to incur strong fitness costs, those that do are recurrently lost in the absence of selection, with the genome undergoing reversion to a single-copy state. Remarkably, once the extra gene copies in the amplified region are lost, there is no remnant trace of the prior presence of CNVs, revealing the ephemeral nature of this class of genetic variation.

## Results

We studied 15 unique lineages of *Saccharomyces cerevisiae* to investigate the fate of adaptive gene amplifications upon relaxing the original selection pressure of nutrient limitation in chemostats (**Figure 1A**). Nine strains (*G_1* - *G_9*) had acquired segmental CNVs containing the *GAP1* gene on chromosome XI in glutamine-limited chemostats. Strain *M_10* has a segmental amplification of *MEP2* on chromosome XIV, which arose during selection in a chemostat fluctuating between glutamine- and ammonium-limitation. Strain *P_11* acquired a segmental amplification containing *PUT4* on chromosome XV during selection in a chemostat fluctuating between glutamine- and proline-limitation. Strains *GM_12* - *GM_15* had acquired whole-chromosome duplications, or aneuploidies, of both chromosomes XI and XIV during fluctuating glutamine- and ammonium-limitation. Using DNA content staining (**Supplemental Figures S1, S2**), we found that four of the 15 CNV strains (*G_1, G_4, G_6, G_9*) are diploid, whereas the other 11 strains are haploid. All diploid strains have identical copies of each chromosome, likely due to autodiploidization following the CNV event [Tung et al. 2021]. To enable direct comparison between haploid and diploid strains, we report ploidy-normalized copy number (PNCN), defined as the absolute copy number of a locus divided by the total number of chromosomal copies in the strain (i.e., one for haploids, two for diploids), throughout the manuscript. These 15 CNV lineages were chosen to survey a range of different features including the driver gene under selection, copy number increase, size of the CNV, molecular mechanism underlying CNV formation, and number of elapsed generations of selection before the lineage was isolated (**Figure 1B** and **Supplemental Table 1**). CNVs contain one of three driver genes (*GAP1*, *PUT4*, and *MEP2*), vary between a copy number of 2 - 4, and span in size from 6 kb to an entire chromosome. The segmental CNVs (*G_1* - *P_11*) were formed by either non-allelic homologous recombination (NAHR), origin-dependent inverted repeat amplification (ODIRA), or transposon insertion followed by NAHR. Aneuploidies were formed by missegregation of the entire chromosome XI, XIV or both. All strains contained CNV reporters that enabled single cell estimation of copy number at the specific locus using flow cytometry. *GAP1* CNV containing lineages contain an mCitrine reporter adjacent to the *GAP1* locus. Additionally, strains *M_10* and *GM_12 - GM_15* have an mCherry reporter next to *MEP2*, and *P_11* has an mCherry reporter next to *PUT4*. Strains *G_4 - GM_15* were obtained from lineage tracking experiments and therefore contain unique barcode sequences enabling sequence-based verification of parental lineages and their descendants (**Supplemental Table 1**). Our study was designed to minimize the potentially confounding effect of other variation– we studied strains that had been isolated after 60-250 generations of evolution, since we expect few point mutations to accumulate in this relatively short timescale. The rate of generation of CNVs is ten times faster than that of SNVs [Avecilla et al. 2022]. Indeed, we found no high-confidence SNVs in any of the 15 CNV lineages.

**Figure 1.**
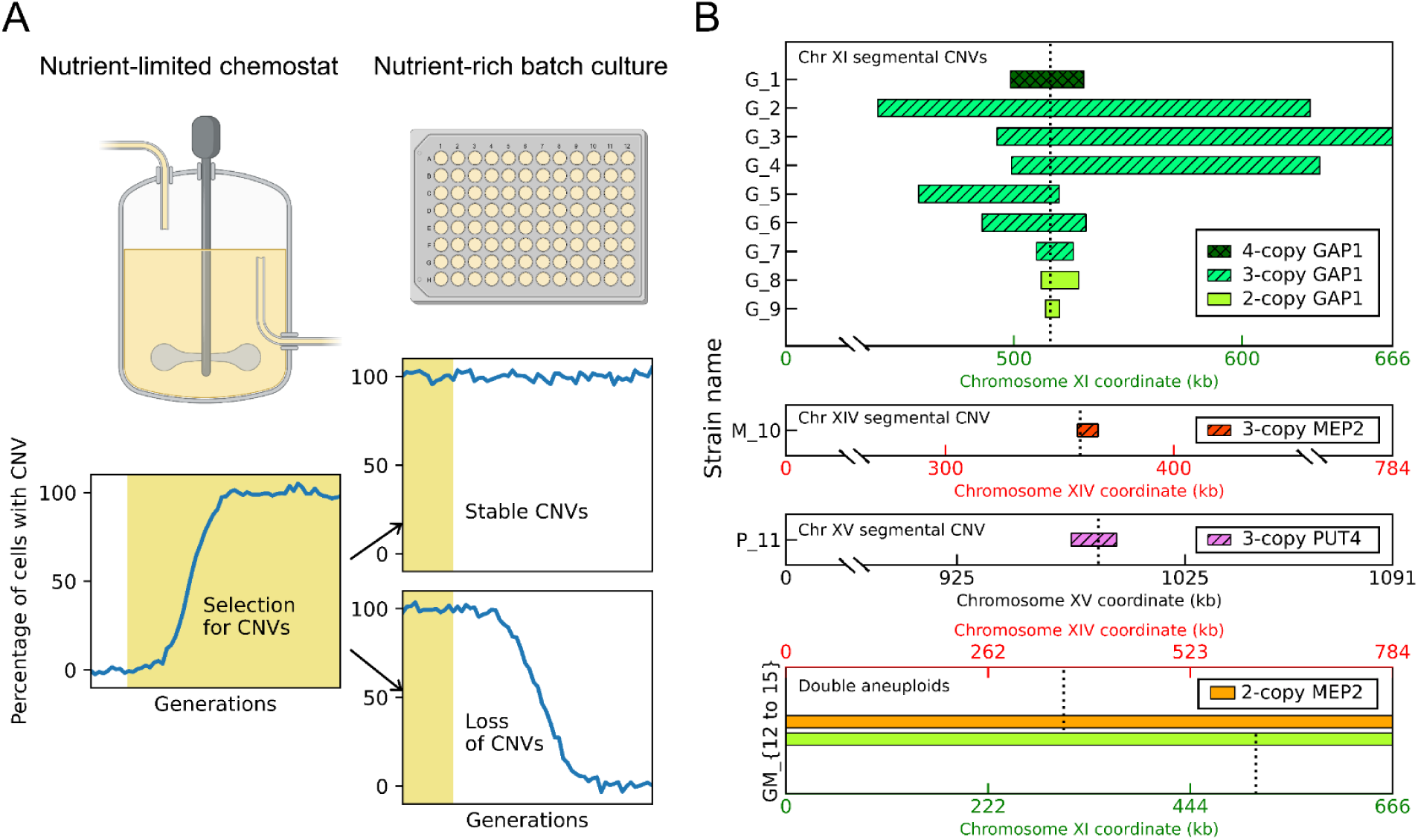
Experimental design. **A)** The two possible fates of adaptive copy number amplifications acquired during nutrient-limited selection in chemostats (yellow shaded background). In the absence of the original nutrient-limitation selection pressure during propagation in nutrient-rich media using 96-well serial batch transfer, if fitness costs are comparable to the benefits, CNVs will be stably maintained (upper panel) whereas if fitness defects of CNVs outweigh their benefits, they will be purged from the population through positive selection for revertants (lower panel). **B)** Schematic representation of the 15 unique CNV lineages used in this study. Our naming convention incorporates three parameters. The gene under selection during amplification is indicated by the letter: *G* (*GAP1* on chromosome XI), *M* (*MEP2* on chromosome XIV), *P* (*PUT4* on chromosome XV) or *GM* (both *GAP1* and *MEP2*). Strains of each type are arranged in descending order of ploidy-normalized copy number (shown by color and shading) and within each copy number, by decreasing size of the amplified region (indicated by length on the x-axis). Positions of the driver genes (*GAP1, MEP2,* and *PUT4*) are indicated by dotted lines.

### Segmental amplifications confer minimal fitness costs in the absence of selection

We sought to test whether adaptive CNVs confer a fitness cost in absence of selective pressure. Therefore, we estimated the fitness of all CNV strains in nutrient-rich media using pairwise fitness competitions against the ancestor (**Figure 2A**). We measured the relative abundance of each CNV strain every 3-4 generations using a flow cytometer and performed linear regression to estimate the per generation growth rate difference, which we define as fitness (**Methods**). We performed all assays in triplicate using a minimum of six time points to maximize our ability to detect small fitness differences.. Interestingly, among the 11 segmental CNVs, only two (*G_4* and *P_11*) have significantly negative fitness effects (p-value < 0.05) in rich media. Surprisingly, two strains containing segmental CNVs showed significantly increased fitness suggesting pleiotropic effects. By contrast, all four double aneuploid (*GM_12 - GM_15*) strains showed significant fitness defects (**Figure 2B**).

**Figure 2.**
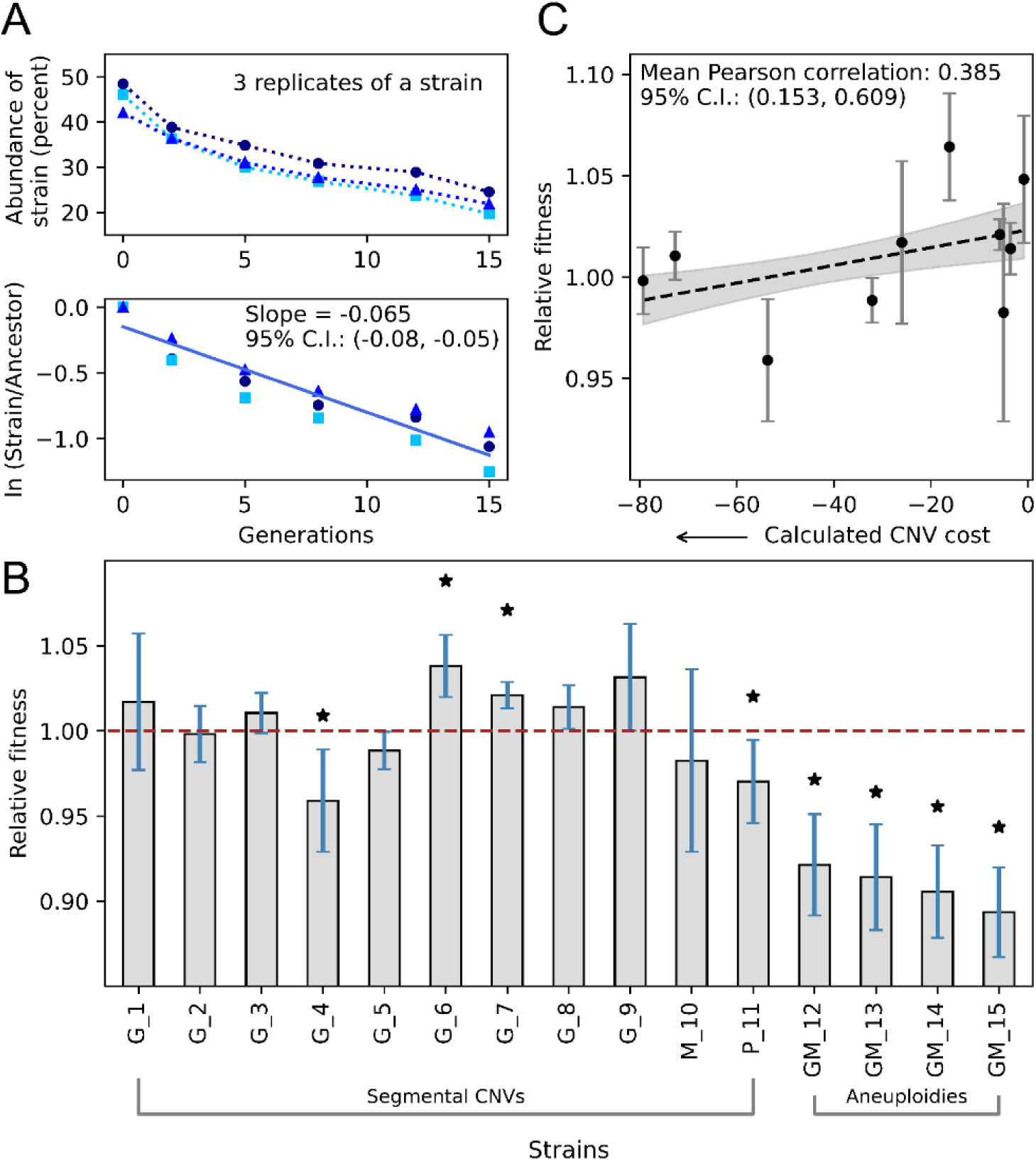
Fitness effects of CNVs. **A)** Pairwise competitive fitness assays were performed in nutrient-rich media by co-culturing each CNV strain with the ancestral single-copy strain and quantifying their relative abundance at multiple time points using flow cytometry (upper panel). Relative fitness was calculated by regressing the natural log of the ratio of the CNV strain and the reference strain against the number of generations, using all 3 replicates for each strain (lower panel). The different shades of blue indicate the 3 independent replicate experiments conducted per strain. **B)** Fitness of all 15 CNV lineages relative to the single-copy ancestor (whose fitness is 1 by definition, marked by a dashed brown line). Each bar shows the fitness of each strain, with error bars indicating 95% confidence intervals from the standard error of the regression using 3 replicates. Strains with fitness significantly different from the ancestor are marked with an asterisk. **C)** The Pearson correlation coefficient was calculated between relative fitness and the estimated fitness costs of segmental CNVs *G_1* - *M_10*. Points with error bars indicate 95% confidence intervals for relative fitness (as in B). Black dashed line and grey shadow indicate 95% confidence intervals for a linear regression through all points, which is shown for visualization purposes.

To investigate determinants of CNV strain fitness, we used data from Rojas et al. [2024], in which the cost of duplicating each individual gene was estimated. We calculated the total cost of each amplified region by summing the individual costs of all significantly positive or negative genes and accounted for non-significant effects using the genome-wide average gene cost (details in Methods). To compute the overall cost of each segmental CNV, the total cost was multiplied by the extra number of copies of the amplified region. Fitness costs were estimated for all segmental CNVs except strain *P_11*, because the fitness cost of several genes within its amplified region were not available in the data. The estimated fitness cost of segmental CNVs is positively correlated (Pearson correlation coefficient, r =0.385; 95% confidence intervals: 0.153, 0.609) with their relative fitness measured using pairwise competition assays (**Figure 2C**). By contrast, the relative fitness of segmental CNVs is not significantly correlated (r=-0.249, 95% confidence intervals: −0.519, 0.024) with the total amount of amplified DNA (**Supplemental Figure S3A**). Together, these results suggest that the fitness cost of a segmental CNV is a joint function of the number of amplified genes and the individual effects of those genes.

### Segmental CNVs are stable in the absence of selection whereas aneuploidies are lost

To investigate CNV stability upon removing nutrient-limited chemostat selection, we performed long-term experimental evolution in nutrient-rich media using serial transfer batch culture. We performed evolution experiments in two sets: 1) strains *G_1* - *G_3* were propagated for 110 generations, and then 2) strains *G_4* - *GM_15* were propagated for 220 generations. Strains *G_1* - *G_3* were evolved in a pilot experiment for 110 generations because in the study where these CNVs arose [Lauer et al. 2018], populations started undergoing changes in copy number well before generation 100. We expect that any loss in copy number would also start occurring on a similar timescale. We observed no loss of CNVs among these three strains by generation 110, indicating their stability. All other strains were subsequently evolved for 220 generations because these CNVs had not only arisen but also attained fixation in their original parent population by generation 200. Hence, observing CNV dynamics till generation 220 was considered a reliable indicator of their stability. We studied 3-4 independent populations per strain. In addition, we simultaneously propagated 5 control strains: (i) S288c unmodified by any reporter gene, which served as a zero-copy control; (ii) mCitrine_1copy, containing a single copy of the mCitrine reporter at the neutral *HO* locus; (iii) mCitrine_2copy, with the reporter at two neutral loci– *HO* and the dubious ORF *YLR123C*; (iv) mCherry_1copy, with a single copy of the reporter replacing the region from the beginning of locus *YLR122C* to the end of locus *YLR123C*; and (v) mCherry_2copy, with the reporter at the *YLR122-123C* and *HO* loci. We confirmed that there was no change in fluorescence in any of these control strains throughout the experiment (**Figure 3A**).

**Figure 3.**
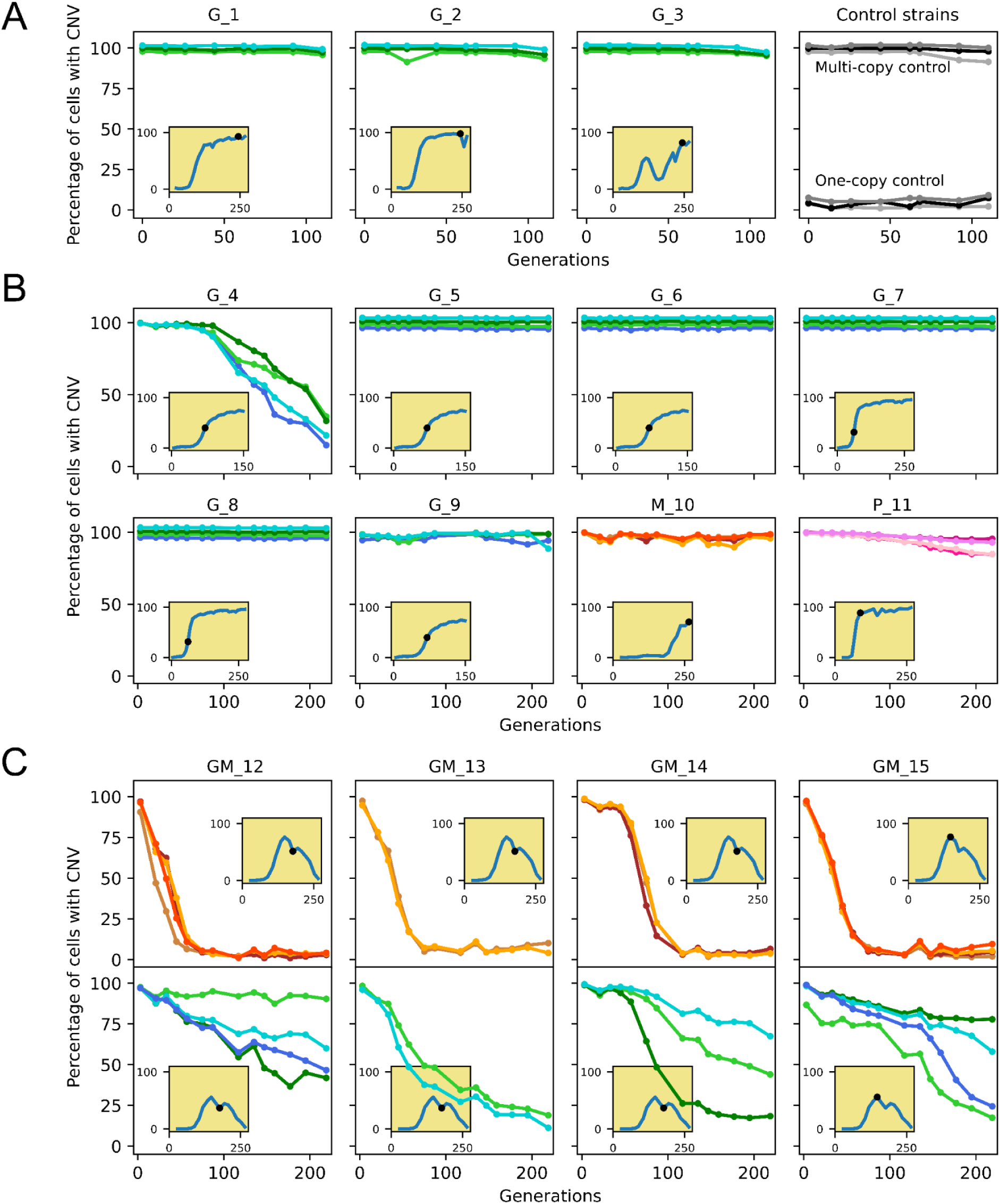
CNV dynamics during long-term experimental evolution in nutrient-rich batch cultures. We quantified the percentage of CNV-containing cells in each population using a fluorescent reporter gene adjacent to each gene of interest. CNVs on Chr XI, XIV and XV, containing *GAP1*, *MEP2* and *PUT4* are shown in green, orange and pink, respectively– different shades indicate independent populations of each strain. Insets show CNV dynamics for each strain’s parental population in which it acquired a gene amplification during nutrient-limited chemostat selection. The timepoint at which the CNV strain was isolated is marked with a dot. **A)** Strains *G_1 - G_3* were evolved for 110 generations. Multiple control strains were propagated that are stable throughout the selection and serve as controls for flow cytometry analysis. **B)** Strains *G_4 - P_11* were evolved for 220 generations. For all strains except *G_4* and *P_11*, data points for all replicates overlap, and therefore have been slightly shifted for better visualization. **C)** Dynamics for both *MEP2* and *GAP1* for double aneuploid strains *GM_12 - GM_15* which were evolved for 220 generations.

Using real time tracking of CNV dynamics we found that 10 of the 11 segmental CNVs were maintained at greater than 95% frequency over 110 (**Figure 3A**) to 220 (**Figure 3B**) generations. There is a small degree of overlap in the fluorescence distributions for one and two copies of reporters, as well as slight fluctuation in fluorescence from one timepoint to another. Consequently, the decrease in fluorescence observed in these 10 strains (e.g. *G_9* and *P_11*) is comparable to the range of fluctuation observed in control strains (**Figure 3A**), and therefore does not reflect reversion of CNVs. Of the 11 segmental amplifications, only one (*G_4*) showed evidence for reversion, which was observed in all independent populations. This indicates that most segmental amplifications that are positively selected in nutrient-limited chemostats are stably maintained upon relaxation of the selective pressure. We find this to be the case for segmental CNVs on chromosome XI, XIV or XV, suggesting that these results are independent of locus and chromosome.

Aneuploidies were lost far more rapidly and consistently than segmental CNVs. All double aneuploid strains (*GM_12* - *GM_15*) exhibited some degree of CNV loss in all but one of the independent populations. In these strains, the chromosome XIV aneuploidy was rapidly lost in all 13 populations whereas the chromosome XI aneuploidy showed variable evolutionary dynamics– some populations lost the chromosome XI faster than the segmental amplification in *G_4*, whereas others maintained it stably for all 220 generations (**Figure 3C**).

### CNV reversion and selection dynamics differ between chromosomes

To quantify the dynamics of CNV loss, we defined three parameters: (i) early phase: the generation at which 25% of cells in a population lost the CNV (ii) middle phase: the generation by which 50% of cells lost the CNV (iii) and late phase: the generation by which 75% of cells lost the CNV (**Figure 4A**). We calculated the median number of generations needed for CNV loss, and performed paired t-tests to compare CNV types in the early phase (**Figure 4B**), middle phase (**Figure 4C**) and late phase (**Figure 4D**).

**Figure 4.**
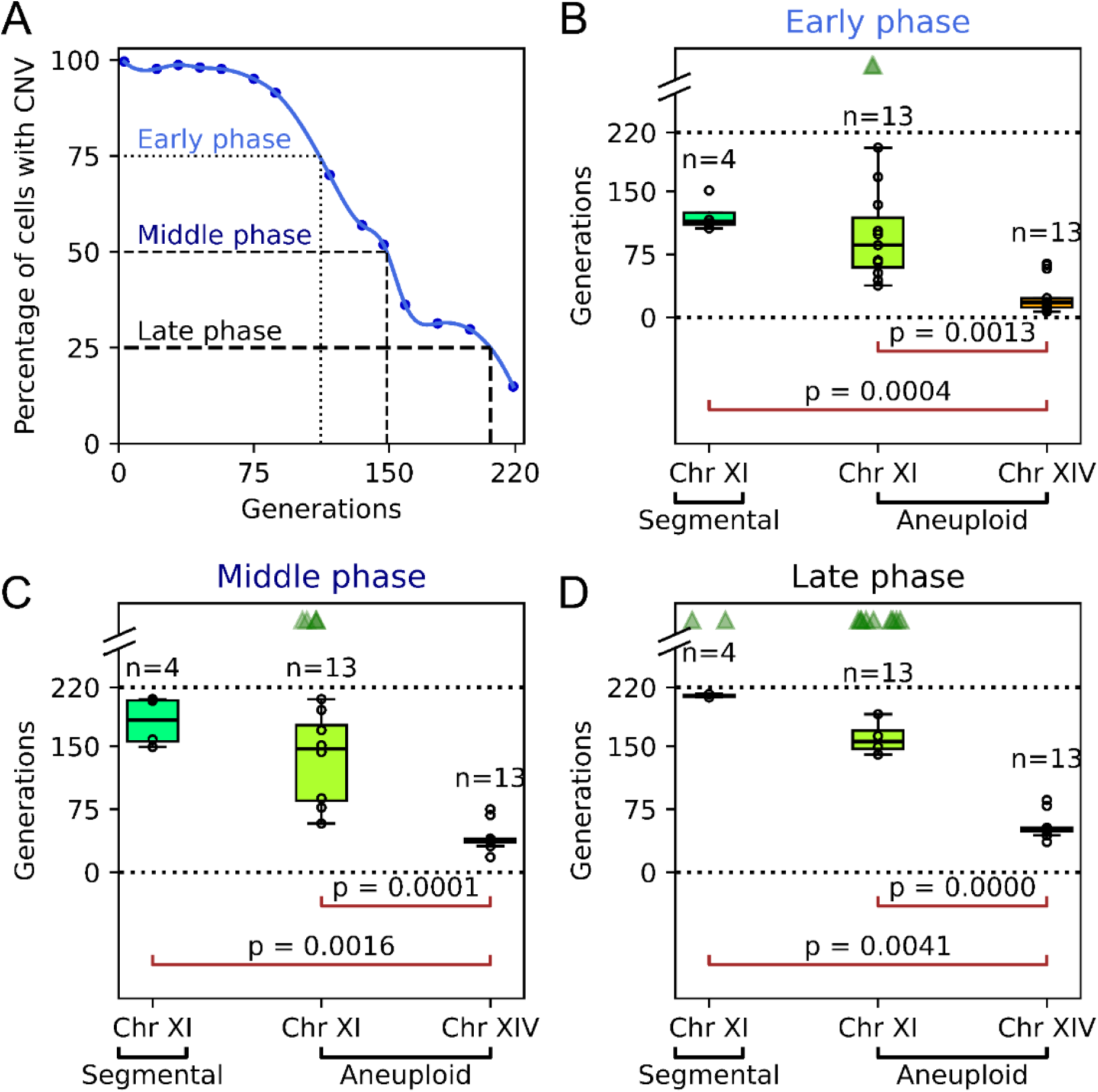
Quantification of CNV loss dynamics. **A)** We define three phases– early, middle and late, corresponding to loss of CNVs in 25%, 50% and 75% cells of each population. **B)** Number of generations to CNV loss and pairwise t-tests between CNV types in the early phase. **C)** Number of generations to CNV loss and pairwise t-tests between CNV types in the middle phase. **D)** Number of generations to CNV loss and pairwise t-tests between CNV types in the late phase. In each phase, sample sizes (n) for each CNV type are shown. CNVs which were lost (*G_4*, *GM_12* - *GM_15*) were grouped by type to perform paired t-tests. P-values are shown for only those pairs which had significant differences. Populations which did not lose the requisite percentage of CNV-containing cells for each phase were excluded from the barplot (but not from the t-test) and are indicated by jittered triangles at the top. Dotted lines indicate the start and end points of the evolution experiment (0 and 220 generations). For each phase, there were significant differences in the dynamics for chromosome XIV aneuploidy (784 kb, 2 copies) versus both the chromosome XI aneuploidy (666 kb, 2 copies) and the segmental amplification of chromosome XI (135 kb, 3 copies), but not between the two kinds of chromosome XI CNVs.

The slowest loss was observed in the chromosome XI segmental amplification, followed by the chromosome XI aneuploidy and then the chromosome XIV aneuploidy, which had the fastest CNV loss. Statistical comparison confirmed that dynamics of CNV loss vary significantly between chromosome XI and chromosome XIV. However, there was no significant difference between the segmental CNV on chromosome XI and the aneuploidy of chromosome XI (**Figure 4**). This was in spite of these two types of CNVs differing in (i) size of the amplified genomic region (135 kb vs 666 kb, respectively), and hence, the number of amplified genes; and (ii) copy number relative to the ancestor (3 versus 2). We note that all strains with chromosome XI aneuploidies also have chromosome XIV duplications. However, since the evolutionary dynamics of the two aneuploid chromosomes are significantly different, we consider them sufficiently independent to be studied separately.

Double aneuploid strains *GM_12* - *GM_15* enable direct comparison of the stability of two different whole-chromosome duplications in the same cell. First, we observed that the extra chromosome XIV was lost very consistently, with all 13 populations across 4 strains showing very similar dynamics, whereas the evolutionary dynamics of the extra chromosome XI were highly variable, both within and across these strains. For instance, at the end of 220 generations, the percentage of cells that lost the extra chromosome XIV ranged between 90-98% across all 13 populations; for chromosome XI, this was between 9-89% for the same populations (**Figure 3C**). Second, the chromosome XIV aneuploidy was lost much faster than the chromosome XI aneuploidy: the median time to early, middle and late phases for populations that lost the CNVs was 18, 39 and 51 generations for chromosome XIV, versus 80, 143 and 149 generations for chromosome XI (**Figure 4**). Finally, analysis of clones from evolved populations after 159 and 201 generations did not identify any lineages that had lost only the extra copy of chromosome XI but not of chromosome XIV. These results indicate that the fitness costs and stability of aneuploidies differ between chromosomes.

### Fitness costs predict CNV reversion in the absence of selection

The five strains that underwent recurrent CNV loss in independent populations (*G_4* and *GM_12* - *GM_15*) were the only ones among our 15 CNV lineages that displayed large fitness defects in rich media (**Figure 2B**). One additional strain, *P_11*, had a small fitness defect compared to these five strains, and showed very slow and minimal CNV loss in 2 of 3 populations (**Figure 3B**). Our results indicate that fitness defects are predictive of the fate of adaptive CNVs in a non-selective environment.

We also estimated fitness costs of aneuploidies of chromosomes XI and XIV separately, by adding the costs of duplicating all the genes on each chromosome, which were measured by Rojas et al. [2024]. The cost of double aneuploidy across both chromosomes was estimated by summing the total costs of the two individual whole-chromosomal duplications. We accounted for non-significant effects using the genome-wide average gene cost. Aneuploidy of chromosome XI had a lower predicted fitness cost (−110.51) than that of chromosome XIV (−117.95), which corresponds to a significant difference in median growth rates (95% and 80% of the euploid growth rate, respectively) [Rojas et al. 2024]. This estimated fitness difference is consistent with the rapid loss of chromosome XIV compared to chromosome XI in double aneuploid strains (**Figure 4**).

We tested whether relative fitness of aneuploidies determined by pairwise competition assays is positively correlated with estimated fitness costs as observed for segmental CNVs (**Figure 2C**). We calculated the relative fitness for double aneuploidy by averaging the observed fitness of strains *GM_12 - GM*_*15* (with whole-chromosomal duplications of chromosomes XI and XIV). For single aneuploidy of chromosome XI, we averaged the observed fitness of eight partial revertants (derived from strains *GM_12* - *GM_15)* which lost the extra chromosome XIV and retained duplication of only chromosome XI. Since there were no revertants which lost only the extra chromosome XI, we could not measure relative fitness for single aneuploidy of chromosome XIV. Both the double and single aneuploidies lie on the regression model based on segmental CNVs (**Supplemental Figure S3B**). This suggests that fitness costs of aneuploids are predictable based on the fitness costs of individual genes.

### Evolutionary model and simulation-based inference

The dynamics of CNV loss are determined by the rate of reversion and the associated fitness effect. To infer the rate of reversion, we modelled the CNV loss dynamics of the five strains (*G_4, GM_12* - *GM_15*) using a Wright-Fisher model and neural network simulation-based inference (nnSBI) (**Figure 5A**). For all strains, the collective posterior predictions of CNV loss dynamics qualitatively match the empirical data (**Figure 5B**). In addition, the overall fitness estimates for these five strains are very similar to results from competition assays, further validating our approach. The reversion rate estimates of both *GAP1* and *MEP2* CNVs vary between 10^-5^ to 10^-3^ per cell per generation (**Supplemental Figure S4**). The fitness estimates of *MEP2* CNV relative to the wild type (0.922-0.949), were significantly lower than those of *GAP1* CNVs (0.971-0.998), suggesting that *MEP2* reversion is under stronger positive selection (**Supplemental Figure S4**).

**Figure 5.**
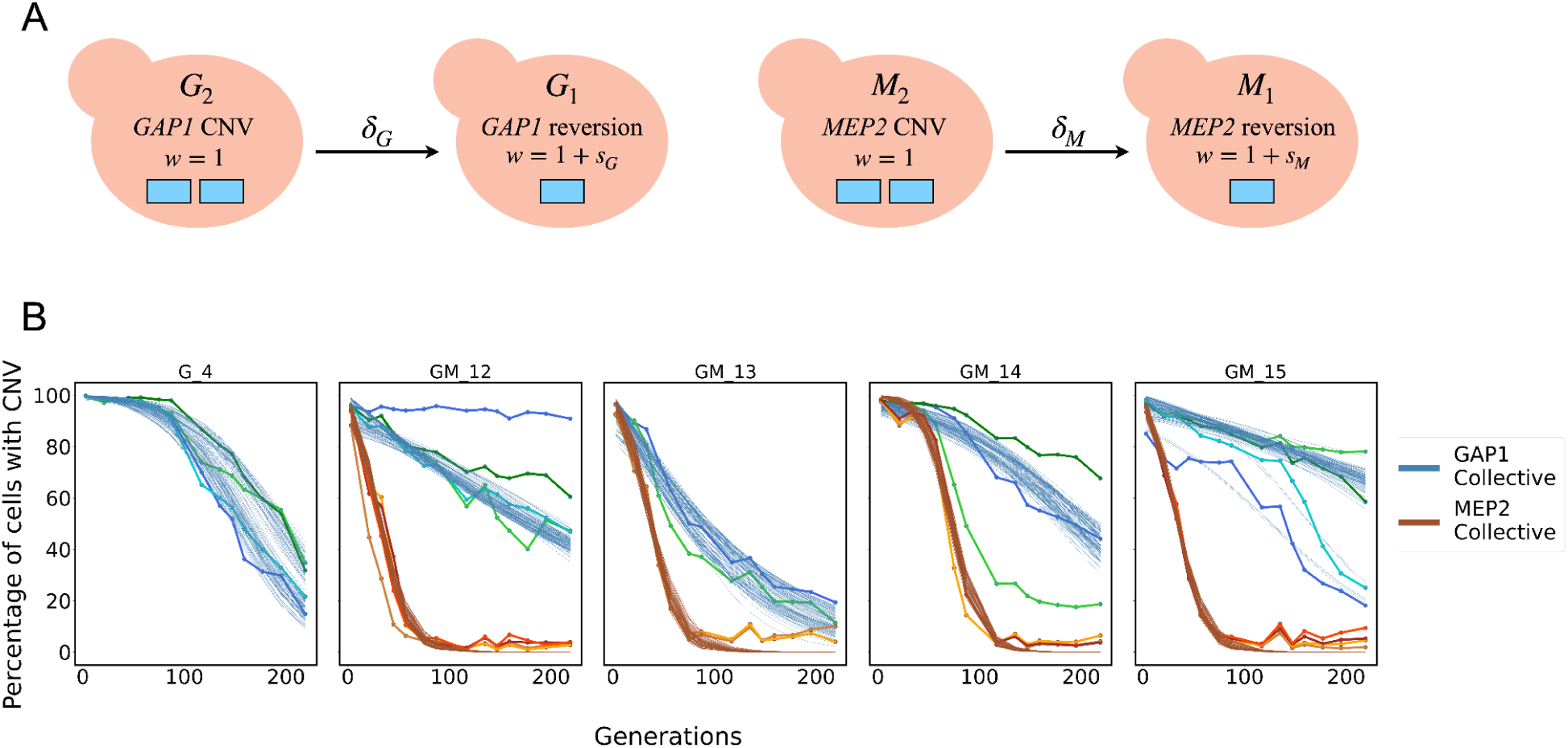
A Wright-Fisher evolutionary model of allele-tracking and collective posterior predictive checks using nnSBI. **A)** CNV reversions are assumed to be independent, with a selection coefficient of *s*_*i*_ and reversion rate of δ_*i*_ for gene *i*. Cell labels refer to a gene (G - *GAP1* or M - *MEP2*) and its copy number (2 or 1) as an allele. **B)** Model simulations of 100 parameter sets sampled from the collective posterior distributions inferred from each experimental replicate (shaded lines) and the empirical data (solid lines).

The individual posterior predictions further support our choice of an evolutionary model without epistasis (**Supplemental Figure S5**). However, we also found that using an evolutionary model with epistasis (**Supplemental Figure S6**) and the same inferred parameters, while assuming independent reversion rates and fitness effects, leads to similar posterior predictions **(Supplemental Figure S7)**. Consequently, we cannot rule out epistasis, but since the simplest model without epistasis explains our results, we assume that epistasis does not play a significant role. Finally, we used the evolutionary model to predict the outcome of short- and long-term experiments in a scenario where reversion is effectively neutral. In this case, we predict the decrease in CNV frequency to be extremely slow, requiring about 22,000 generations before the revertant allele is fixed in the population (**Supplemental Figure S8**).

### CNV reversion increases fitness

To test the effect of CNV reversion on fitness, we performed pairwise competitive fitness assays after evolution for the five strains that underwent CNV loss. From populations evolved for 159 or 201 generations, we isolated clones with either partial or complete loss of CNVs. For the diploid segmental CNV in *G_4*, where the amplified region had 3 copies per copy of chromosome XI, ‘partial’ revertants lost only 1 of the extra copies, whereas ‘complete’ revertants lost both extra copies. For double aneuploid strains *GM_12* - *GM_15*, ‘partial’ revertants lost only the extra chromosome XIV, whereas ‘complete’ revertants lost the extra copy of both chromosomes, XIV and XI.

For all five CNVs, partial or complete reversion led to improved fitness in rich media compared to the CNV strain. Some revertants were slightly less fit than the ancestor, but in most cases, reversion restored fitness to levels comparable to or slightly higher than the ancestor. Surprisingly, complete revertants were not universally fitter than partial revertants of the same strain. For example, for strain *GM_15*, the median fitness coefficient before CNV loss was 0.89, which improved to a range of 0.93 to 1.01 for partial revertants, and from 0.94 to 0.99 for complete revertants (**Figure 6**).

**Figure 6.**
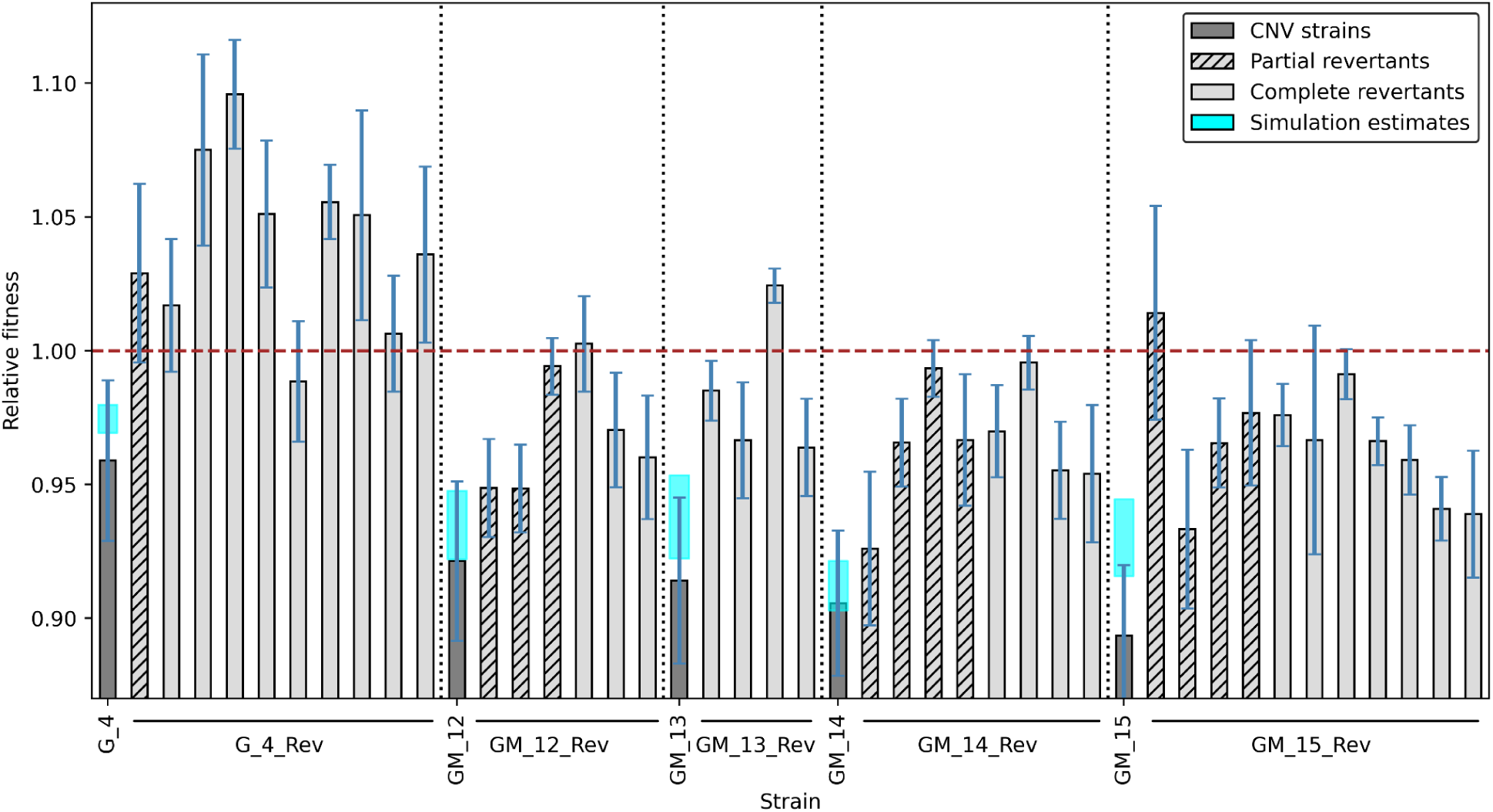
CNV reversion increases fitness. Pairwise competitions were performed to compare fitness before and after evolution in nutrient-rich media. Revertants were isolated after 159 or 201 generations of evolution from populations that underwent CNV loss. For the segmental amplification in *G_4* which has 3 copies, the partial revertant has 2 copies, and complete revertants have 1 copy. For the double aneuploid strains *GM_12* - *GM_15*, partial revertants lost only the extra copy of chromosome XIV, whereas complete revertants lost the extra copy of both chromosome XI and chromosome XIV. Each bar shows the median relative fitness, with error bars indicating 95% confidence intervals calculated from 3 replicates for each strain. Fitness estimates for the CNV strains from our simulation-based inference are similar to our empirical data.

### Single nucleotide variation partly explain fitness differences between similar copy numbers

Although all revertants have higher fitness than their parent CNV strain, we observed some fitness differences between revertants with similar copy numbers. This may be partly explained by certain point mutations that arose over the course of 220 generations of evolution. For example, among complete revertants of *GM_15*, which lost extra copies of both chromosomes XI and XIV, three had much lower fitness than others. Interestingly, these three revertants had a total of four confirmed SNVs, two of which are known to be missense mutations in essential proteins (*GM_15_Rev7*, *GM_15_Rev8*, *GM_15_Rev12*, see **Supplemental Table 5**). Again, among partial revertants (which lost only the extra chromosome XIV) of *GM_15*, the one with the lowest fitness (*GM_15_Rev6*) was found to have a missense mutation. Not all fitness differences between similar-copy revertants can be explained by point mutations– this may be because these slight fitness differences are simply not selected against, as long as they are more fit than the CNV parent.

### CNV reversion can be scarless

To study the genomes of strains after CNV reversion, we isolated a total of 56 clones from 17 revertant populations derived from five strains (segmental *G_4*, aneuploid *GM_12* - *GM_15*) at generation 159 or 201, and performed whole-genome sequencing. We obtained the ploidy-normalized copy number (PNCN) of all genes, relative to the single-copy ancestor, by calculating sequencing depth for each nucleotide normalized to the genome-wide mean, and then averaged over all copies of the chromosome (i.e., 1 for haploid, 2 for diploid) (**Methods**). We confirmed that for all clones that showed evidence of CNV reversion on the basis of the fluorescent reporter, the entire CNV region had reverted to a single copy. For the only segmental amplification that was lost (strain *G_4*), we analyzed sequencing read depth for 15 clones from 4 independently evolved populations (**Supplemental Figure S9**). One clone was a partial revertant, having reverted from a PNCN of 3 copies of the amplified region in the CNV parent to 2 copies, whereas all other 14 clones were complete revertants with a single copy of all genes.

To further investigate cases of partial and complete reversion, we performed long-read sequencing of the CNV parent *G_4*, the partial revertant, and four complete revertants, one from each independent population. Read depth from long-read sequencing had very little noise, confirming that the parental CNV strain had a PNCN of 3 across the entire amplified region (**Figure 7A, Supplemental Figure S11A**). We confirmed that the copy number of revertants inferred from fluorescence corresponded to that obtained from relative read depth (**Figure 7B**). We extracted split reads to find mismatches with the reference genome, which indicate the breakpoints, or boundaries, of the CNV. Both long- and short-read sequencing confirmed that the parental CNV strain had 2 peaks of split reads corresponding to the two breakpoints of the amplified region of the genome (red lines, **Figure 7A, Supplemental Figure S9**).

**Figure 7.**
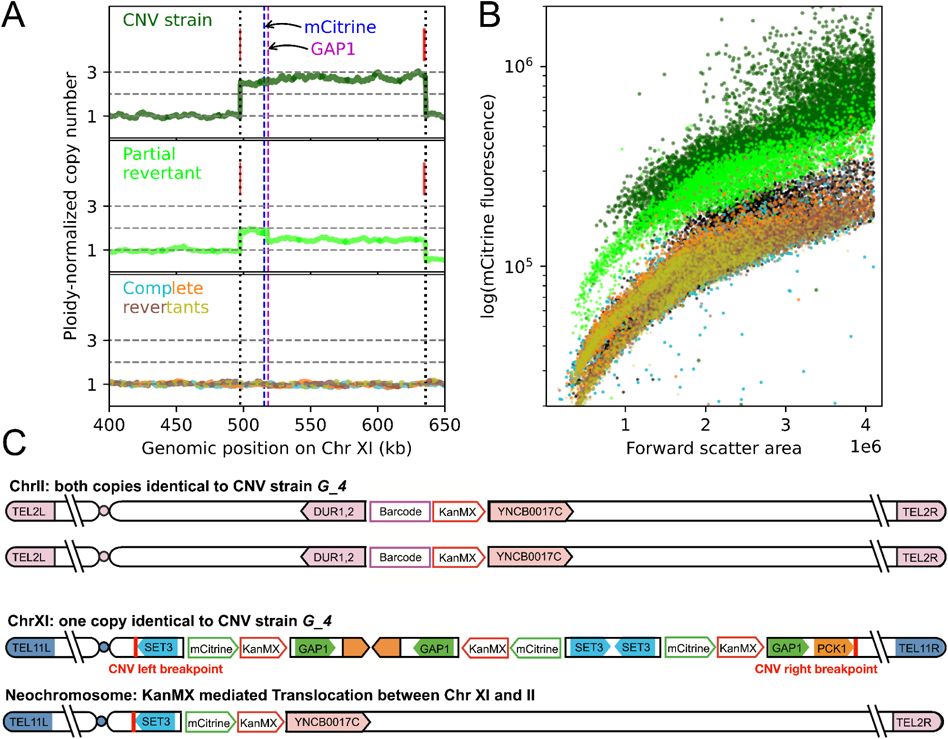
Long-read sequencing and *de novo* genome assembly confirm scarless reversion and reveal mechanism of partial reversion. A) Sequencing read depth for each nucleotide normalized to the genome-wide average, and averaged over two copies of the chromosome for diploids, is the ploidy-normalized copy number (PNCN). Split reads (red lines), found where the sequence differs from the ancestral reference, indicate CNV breakpoints. CNV boundaries are marked by vertical dotted black lines; horizontal dashed grey lines indicate PNCN of 1, 2 and 3 for visual comparison. Vertical dashed lines indicate the positions of *GAP1* (magenta) and its reporter mCitrine (blue). Shown here are the segmental CNV that underwent reversion (*G_4*), a partial revertant, and four independently evolved complete revertants (overlaid) – all diploid. **B)** Fluorescence data indicates PNCN of 3, 2 and 1 corresponding to the CNV strain, partial revertant and four complete revertants, respectively. All six strains are shown using the same colours as in (A). Additionally, the ancestral haploid strain (PNCN =1) is plotted in black. **C)** Partial reversion in copy number occurred due to translocation between the barcode-associated kanMX locus on Chr II, and the leftmost copy of the KanMX associated with the mCitrine reporter locus on Chr XI. The partial revertant strain retains both parental copies of Chr II, but has one copy of the original Chr XI and one copy of the neochromosome produced by the translocation. CNV breakpoints, adjacent to the *SET3* and *PCK1* genes, are marked with red lines.

The partial revertant retained split reads at the same positions as the CNV parent. This evidence, along with the decrease in fluorescence to a level between the 3-copy CNV parent and the 1-copy complete revertants (**Figure 7B**), suggested that the partial reversion had occurred by loss of only one copy of the amplified region. However, the long-read data enabled resolution of the amplified region into two distinct parts– a region with a PNCN of 2 followed by a region with a copy number of 1.5. The unamplified region to the right of the CNV also had a decrease in PNCN from 1 to 0.5. In addition to these changes on chromosome XI, a region on chromosome II had an increase in PNCN from 1 to 1.4, exclusively in the partial revertant (**Supplemental Figure S10, S11A**). The relative read depth of these three regions (red dashed rectangles, **Supplemental Figure S11A)** substantially deviated from the single copy loss model assumed based on previous evidence. DNA staining showed that like the CNV parent, the partial revertant was diploid (**Supplemental Figure S2**).

To define the mechanism of partial reversion, we extracted split reads around identified breakpoints and performed *de novo* genome assembly to create contigs. The contig comprising the left side of the fluorescent CNV reporter (mCitrine) on chromosome XI and the right side of the barcode site (*DUR 1,2*) on chromosome II, suggested a translocation between the two chromosomes, occurring at the KanMX locus (**Supplemental Figure S11B**). Manual verification using select reads that spanned both chromosomes found that these reads were present in both contigs. The translocation resulted in a neochromosome featuring the region of Chr XI ranging from the left telomere (TEL11) to the mCitrine reporter, followed by the region of Chr II from the YNCB0017C tRNA to the right telomere (TEL2R) (**Figure 7C**). The other copy of chromosome XI remained identical to that of the CNV parent. This translocation-mediated mechanism, resulting in two non-identical copies of chromosome XI, is consistent with the copy number across the CNV region (**Supplemental Figure S11A**). It also resulted in a total of four mCitrine reporter copies across two copies of chromosome XI, producing fluorescence corresponding to a 2-copy average (**Figure 7B**).

Complete revertants had a uniform PNCN of 1 across the entire genome (**Figure 7A**). None of the 14 complete revertants were found to retain any split reads via long- or short- read sequencing, confirming that complete loss of the triplicated region occurred in all populations. There were also no rearrangements or translocations anywhere in the genomic sequence outside the previously amplified region. This suggests that the reversion to the one-copy genome does not leave any sequence evidence of the prior presence of gene amplifications. When we compared the nucleotide sequence in the breakpoint regions, we found that the inversion present in the CNV strain is fully reversed in all complete revertants, making the revertant sequence indistinguishable from the reference (**Supplemental Figure S12**). These complete revertants were verified to be diploid (**Supplemental Figure S2**), indicating that both copies of chromosome XI in complete revertants lost the entire amplified region).

We propose a mechanistic model that likely underlies complete reversion of the segmental 3-copy CNV in strain *G_4* (**Figure 8**). The CNV is a triplication on chromosome XI, with the middle copy (green) inverted with respect to the ancestral genome (**Figure 8A**). Since the first (blue) and third (red) copies are identical to each other and in the same orientation, they can align to result in homologous recombination within one of the copies of chromosome XI. Such a crossover event results in a complete deletion of the middle copy. The recombined genomic sequence is identical to the single-copy ancestral strain prior to CNV amplification. Now the cell is heterozygous for chromosome XI, with one revertant copy identical to the ancestor and the other copy identical to the CNV strain (**Figure 8B**). Following replication during mitosis, duplicated sister chromatids undergo segregation, and identical copies of chromosome XI are sometimes passed down to the same daughter cell. Since the CNV in strain *G_4* confers a fitness defect in the absence of positive selection (**Figure 2B)**, daughter cells having both revertant copies of chromosome XI are likely to have a fitness advantage over cells with two CNV-containing chromosomal copies, and over cells heterozygous for CNV and revertant copies. Thus, complete revertants will ultimately go to fixation, taking over the entire evolving population.

**Figure 8.**
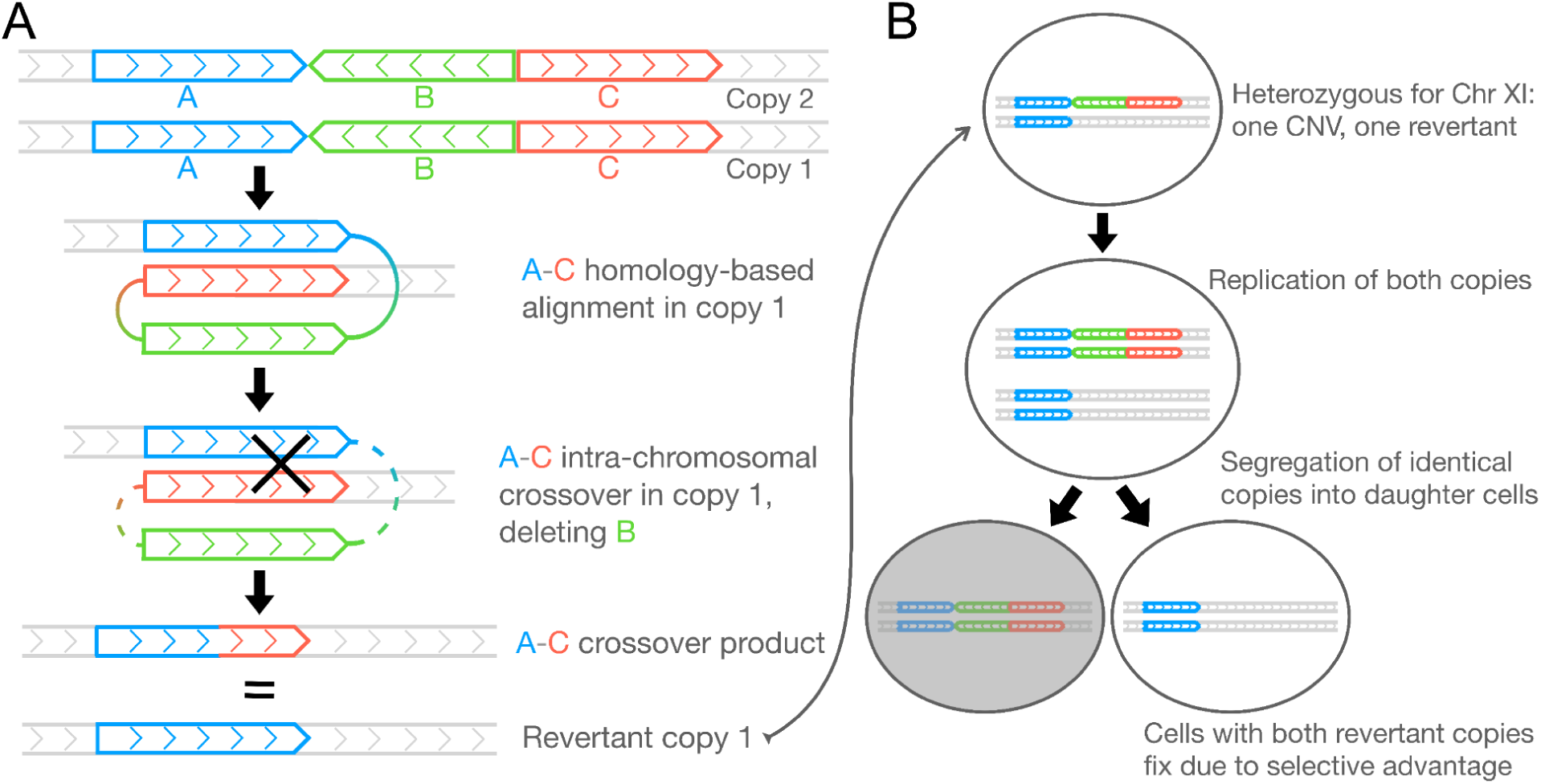
Mechanistic model for complete reversion of the diploid segmental CNV. **A)** Reversion first occurs by intrachromosomal homologous recombination, in one chromosome, generating a chromosome identical to the ancestor. **B)** In the heterozygote, segregation of two identical sister chromatids into one cell during mitotic division creates complete revertants. Cells with both CNV-containing chromosomes have a fitness disadvantage compared to cells with both revertant chromosomes, and are thereby outcompeted in the population.

## Discussion

In this study, we sought to investigate the long-term fate of adaptive gene amplifications in budding yeast upon reversal of the selection pressure in which they arose. By evolving 15 unique CNV lineages in nutrient-rich media, we removed the original selection pressure of nitrogen limitation in chemostats, which had imposed positive selection for *de novo* CNVs containing nutrient transporter genes. We used experimental evolution, flow cytometry, fitness assays, simulation-based inference, and whole-genome sequencing to study the dynamics of these adaptive CNVs when returned to their pre-selection environment.

CNVs often incur fitness costs associated with elevated gene expression and stoichiometric imbalances of encoded products [Tang and Amon 2013, Gerstein and Berman 2015, Tsai et al. 2019, Yang et al. 2021, Rojas et al. 2024, Spealman et al. 2025]. Fitness costs are balanced against fitness benefits conferred by these CNVs, which are, typically, context dependent. We hypothesized that upon removal of the selection pressure imposed by the nitrogen-limited environment, CNVs containing nitrogen-transporter genes would no longer confer a fitness advantage, but fitness costs would remain, resulting in lower fitness relative to the single-gene copy ancestor.

However, contrary to our expectation, only two of our 11 lineages containing segmental CNVs had reduced fitness compared to the ancestor in nutrient-rich media. These results differ from our prior analysis of CNV strains that arose in glutamine-limited chemostats [Avecilla et al. 2023]. We attribute this difference to the difference in conditions in the two studies (Avecilla et al. 2023 used galactose-containing media) and the increased sensitivity of our pairwise fitness assays.

Tracking CNV dynamics using fluorescent CNV reporters showed that in the absence of a fitness cost, most segmental CNVs were stably maintained across hundreds of generations in nutrient-rich conditions. By contrast, segmental CNVs with significant fitness defects in a nutrient-rich environment were lost from evolving populations as there was positive selection for spontaneously generated revertants containing a single copy of the locus. Our results regarding general CNV stability are similar to some previous studies that probed CNVs in yeast under selection conditions different to our study. For example, segmental duplications within a chromosome are stably inherited even in the absence of any selective advantage, as indicated by growth rates comparable to wildtype [Koszul et al. 2006]. Conversely, CNVs arising under antifungal drug treatment in *C. albicans* rapidly return to the progenitor copy number once the selection pressure is relaxed [Todd et al. 2020]. Our result that most CNVs are stable also differs from previous studies in other organisms such as *Salmonella* and *E.coli* [Reams et al. 2010, Tomanek et al. 2020], which showed that due to high cost and high rate of deletion, copy number amplifications often return to the ancestral single-copy state in the absence of selection. Conversely, our findings are consistent with the observation that copy number variation at the *CUP1* locus is maintained among commonly used lab strains of *S. cerevisiae*, suggesting that these CNVs are maintained despite the absence of continued positive selection [Warringer et al. 2011]. Overall, we conclude that fitness costs of CNVs in absence of the original selection pressure are variable and context-dependent, likely depending on the specific environment, genetic background, and CNV structure. When a CNV does incur a significant fitness defect in the non-selective environment they revert and are rapidly purged from the population.

In contrast to segmental CNVs, all four strains with aneuploidies of chromosomes containing nitrogen transporter genes had lower fitness relative to the ancestor in the non-selective environment. These aneuploid lineages were positively selected in nitrogen-limited environments, but show clear fitness defects in rich media. Previous studies have found that engineered aneuploids grow more slowly than euploids, regardless of karyotype [Torres et al. 2007; Sheltzer et al. 2012; Beach et al. 2017] and adaptive aneuploids can also show antagonistic pleiotropy such that they are deleterious in other environments [Sunshine et al. 2015; Linder et al. 2017]. Aneuploidies were lost more frequently than segmental CNVs among our lineages, which agrees with previous studies that showed that large duplications on supernumerary chromosomes and disomic chromosomes are highly unstable and are spontaneously lost at high frequency [Campbell et al. 1975, Koszul et al. 2006]. Aneuploidy can serve as a fast but transient evolutionary solution during adaptation to a selection pressure [Yona et al. 2012, Kohanovski et al. 2024]. Our results are consistent with the instability and dynamic nature of aneuploidies.

As the four aneuploid lineages contained two copies of both chromosome XI and chromosome XIV, these strains provided an opportunity to directly compare the stability of two different chromosomes. We found that all evolving populations purged the extra copy of chromosome XIV much faster and more ubiquitously than the extra copy of chromosome XI. This suggests that aneuploidy of chromosome XI has a lower fitness cost than that of chromosome XIV, and hence, is better tolerated. This result is in agreement with Rojas et al. [2024]. Thus, the differing fitness costs of chromosomal aneuploidies are predictive of their long-term stability. Simulation-based inference suggests that reversion rates are similar for the two chromosomes, but the fitness cost associated with chromosome XIV aneuploidy is higher. The estimated reversion rates are higher than the CNV formation rates previously reported [Chuong et al. 2025; Avecilla et al. 2022], which had aggregated both aneuploidies and the less frequent focal CNVs. The fitness costs and rapid reversion rate indicates that spontaneous aneuploidies are expected to be rapidly lost from populations in the absence of positive selection.

Our study design allowed direct comparison of aneuploidy of chromosome XI and segmental CNVs on chromosome XI. We observed that aneuploidies of chromosome XI experienced negative selection in the vast majority of cases (12 out of 13 populations), whereas segmental CNVs in chromosome XI were lost rarely (4 out of 33 populations, or 1 of 9 unique segmental CNV lineages). We did not find a correlation between fitness costs of CNVs and the number of additional bases in the CNV region, consistent with our prior findings [Avecilla et al. 2023]. The sole segmental CNV that repeatedly reverted had no obvious distinguishing features compared with other segmental CNVs that were maintained during the evolution experiments. This suggests that fitness costs are not fully explained either by gene load or by specific dosage-sensitive genes, as concluded by Rojas et al. [2024].

Analysis of the sole segmental CNV that underwent reversion and selection revealed a perfect reversion to the ancestral single-copy genome. We confirmed that reversion was scarless using both short-read Illumina sequencing and long-read PacBio sequencing data of several independent isolates. Our finding is supported by previous studies across different species. In *C.albicans*, antifungal drug resistance can drive CNVs that are reversible upon relaxation of this selection pressure, and can leave no trace that the CNV ever existed [Todd et al. 2020]. In *S.cerevisiae*, scarless reversion has been seen in independent instances of loss of *CUP1* amplifications [Chang et al. 2013]. In *E.coli*, gene amplifications can act as a fast but transient response that disappears once environmental pressure is relaxed, leaving no genomic signature [Tomanek et al. 2020, Tomanek et al. 2022]. Our results, together with existing literature, provide strong evidence that complete CNV reversion might be a common phenomenon underlying rapid adaptation in dynamic environments.

Our results provide new insights into the plasticity and adaptability of eukaryote genomes. The transient and reversible nature of CNVs makes them an ephemeral adaptive class of genetic variation that may be a more widespread driver of adaptation in natural populations than inferred based on their observation in extant populations. Our results suggest CNVs can allow organisms to flexibly switch between phenotypes as is beneficial, allowing rapid adaptation to fluctuating environments. One limitation of this study is that all CNVs were nutrient transporter-driven amplifications that arose during evolution in nutrient limitation in chemostats. Although our CNV lineages include unique nutrient transporter loci on three different chromosomes, it would be interesting to study the stability and reversibility of adaptive CNVs originating in other kinds of environments since fitness costs vary based on environmental and genetic context, as discussed earlier.

## Materials and Methods

### Strains and media

We used 15 budding yeast strains with genome amplifications evolved from the haploid derivative of the *Saccharomyces cerevisiae* lab strain S288c. This strain was first modified to have an mCitrine gene under the control of the constitutively expressed *ACT1* promoter (ACT1pr::mCitrine::ADH1term) and marked by the KanMX G418-resistance cassette (TEFpr::KanMX::TEFterm). This mCitrine CNV reporter construct is on chromosome XI, adjacent to the *GAP1* gene. Strains *G_1* - *G_9* were isolated from this modified ancestor during a prior LTEE in glutamine-limited chemostats, described in Lauer et al. (2018). Cells from evolved populations of the ancestor preserved using glycerol at −80°C, were plated on glutamine-limiting media. Colonies were selected for increased mCitrine fluorescence using blue light, and grown in glutamine-limiting cultures overnight. The fluorescence of CNV-containing clones was validated using the flow cytometer Cytek Aurora and they were then preserved at −80°C using glycerol.

We then further altered the modified ancestor to include an mCherry reporter either on chromosome XIV beside the *MEP2* gene, or on chromosome 15 adjacent to the *PUT4* gene. *MEP2* and *PUT4* were amplified in ammonium-limited and proline-limited chemostats, respectively. Strains *M_10 - GM_15* were evolved from such doubly modified ancestors with two reporters near two different genes of interest (*GAP1* and *MEP2* or *PUT4*) during another previous LTEE (Abdul-Rahman et al., in prep) in which dual-fluorescence strains were evolved in fluctuating conditions of two different nitrogen-limiting sources: either glutamine and ammonium sulfate or glutamine and proline. Populations were revived from glycerol stocks archived at −80°C by inoculating overnight cultures of glutamine-limiting media and incubating them at 30°C. Fluorescence-Activated Cell Sorting (FACS) was used to sort subpopulations based on their FITC-A vs mCherry-A profiles in which all pairwise combinations of mCitrine:mCherry copy numbers ranging from 0 to 3 were gated separately (i.e. 0:0, 0:1, 0:2…) by using the 0-, 1-, and 2-copy controls. For each subpopulation, 100,000 - 200,000 cells were collected into 500 uL of 1X PBS, 100 uL of which was inoculated into an overnight culture of glutamine-limited media. Clones from each overnight culture were then isolated and mCitrine and mCherry copy numbers were validated by performing flow cytometry using the Cytek Aurora and they were then preserved at −80°C using glycerol.

To make fluorescence control strains, the mCitrine or mCherry reporter was incorporated at one or both of these neutral loci: *HO (YDL227C)* on chromosome IV and the dubious ORF *YLR123C* on chromosome XII. All experiments in the present study were performed in optimal growth conditions for laboratory strains of budding yeast using yeast extract peptone dextrose (YPD) media and growth at 30°C.

### Ploidy assay

We measured ploidy of CNV strains and revertants of *G_4* by fixing cells with 70% ethanol in exponential growth phase and staining DNA with propidium iodide. We followed the protocol described in Todd et al. [2018] to prepare stained samples. We incubated samples overnight and measured fluorescence (channel YG4-A) on the Cytek Aurora. Using 3-4 replicates per strain, we inferred ploidy by comparing the fluorescence distribution of each strain to that of haploid and diploid control strains.

### Long-term experimental evolution

We performed experimental evolution of all CNV strains via serial dilution in batch culture in 96-well plates. For each CNV strain, we isolated 3-4 isogenic clones to found independent populations that were placed in randomly-assorted wells. We cultured each clone in YPD, with a total volume of 200 μl per well. We back-diluted the cultures in fresh media by 1:64, such that they went through the full growth cycle of lag phase, exponential phase, and stationary phase every 6 generations. Strain *G_1 - G_3* were evolved for 110 generations, and strains *G_4 - GM_15* were evolved for 220 generations. We sealed the plates with a breathable membrane to prevent cross-contamination between wells while maintaining an aerobic environment at 30°C, shaking at 170 rpm.

### Pairwise fitness competition assays

To measure the relative fitness of CNV strains and revertant clones in rich media, we performed pairwise competitive fitness assays. We co-cultured each strain of interest with the haploid reference strain lacking any fluorescent reporter, for 8-16 generations in conditions identical to the evolution experiment (96-well plate batch culture in YPD at 30°C, shaking at 170 rpm). We measured the relative abundance of each strain via the flow cytometer Cytek Aurora, and plotted the natural log of the ratio of the two strains, every 2-4 generations. This assay was done in triplicate for each strain, so that we could fit a linear regression using all three independent values. The starting ratio for each assay was set to have the same value, making it a one-parameter linear regression where we only fit for the slope. Relative fitness of each strain was calculated as (slope + 1), equal to 1 for the reference strain.

### Estimation of CNV fitness costs

We analyzed the data in GeneDuplication_fitnessCost_and_compiled_features.xlsx (Table S2) of Rojas et al. [2024]. This paper had quantified the fitness effect of duplicating each gene as a log_2_fold change in abundance after competitive growth. We calculated the total cost of each amplified region by adding the individual fitness costs of all significantly positive or negative genes (false discovery rate < 0.05). Similar to the best-fit model in that paper, we also added all non-significant genes located within the CNV by replacing each of their costs with the mean cost of all genes. To get the final cost of each segmental CNV, this total cost was multiplied by the number of extra copies of the amplified region. Similarly, for calculating the cost of an aneuploidy, fitness costs of all genes present on the duplicated chromosome were accounted for.

Total cost of a CNV = Extra copies of amplified region x [Sum of fitness costs of all significantly positive and negative genes + (Number of genes with non-significant costs x Mean cost of all genes)], where Extra copies of amplified region = (Copy number-1) and Mean cost of all genes = −0.33 (from Rojas et al. [2024]).

### Flow cytometry

To monitor CNV dynamics in evolving strains using their fluorescent reporters, we sampled 20ul from each 200ul population and analyzed them on a Cytek Aurora flow cytometer. We measured 10,000 cells per population, recording mCitrine fluorescence signal (excitation = 514 nm, emission = 530 nm, using the B2-A filter), mCherry fluorescence (excitation = 587 nm, emission = 610 nm, using the YG3-A filter), cell size (forward scatter), and cell complexity (side scatter). We sequentially filtered out debris using side scatter area (SSC-A) and doublets using forward scatter height (FSC-H) and forward scatter area (FSC-A). We then measured fluorescence for each cell and divided this by FSC-A to account for differences in cell size. To quantify the percentage of cells in each population that have zero, one, and two or more copies of the genes of interest, we used control strains with known copy numbers of the reporter genes to manually draw gates around the range of normalized fluorescence values that correspond to each copy number. Since there is some overlap between the distribution of values corresponding to one and two copies, the process of cell selection via gating inherently involves some misclassification. We drew the one- and two-copy gates such that they were mutually exclusive, with the fraction of misclassification kept to a minimum. This ensured that each copy number gate contained 95% of cells of the corresponding control strain. The proportion of cells in the ‘2 or more copy’ fluorescence gate at each timepoint allowed us to track the percentage of CNV-containing cells in each population over generations.

Prior to the evolution experiment, we optimized our flow cytometry protocol to ensure highest expression of fluorescent reporter genes, leading to maximal separation between the fluorescence distributions of one copy vs two copies of a reporter, and decreasing errors in gating. Fluorescence signal is reduced when cells are in stationary phase, so flow cytometry should be done in exponential phase. We measured fluorescence at different timepoints after each serial dilution, which switches cells from stationary phase back to exponential growth. We found that fluorescent signals are at their peak 4-5 hours after dilution.

### Isolation of revertant clones from evolved populations

For isolating revertant clones from evolved populations preserved using glycerol at −80°C, we chose two late timepoints, generations 159 and 201, as representative of the latter part of the evolution experiment, by which revertant populations had lost CNVs in all or a majority of cells. We plated cells on YPD from populations that showed CNV loss. Colonies were preliminarily screened for mCitrine fluorescence using blue light, from which randomly selected colonies were grown independently in liquid YPD overnight. We then measured the fluorescence of each culture using the flow cytometer Cytek Aurora. Clones with fluorescence corresponding to one copy of one or both of the reporter genes (mCitrine and mCherry) were designated as revertants. A total of 56 revertant clones from 17 populations across 5 strains were then preserved at −80°C using glycerol, for subsequent sequencing and fitness assays.

### Whole-genome sequencing

For CNV strains and revertant clones isolated from their evolved populations, we extracted genomic DNA using the Hoffman-Winston protocol [Hoffman and Winston, 1987]. We quantified gDNA concentration using dsDNA broad range reagents on the Qubit fluorometer. DNA libraries were sequenced using a paired-end protocol (2 × 150) protocol on an Illumina Nextseq 500 or NovaSeq 6000. The reads were basecalled using Picard IlluminaBasecallsToFastq version 2.23.8 [https://broadinstitute.github.io/picard/], with APPLY_EAMSS_FILTER set to false. Following basecalling, the reads were demultiplexed using Pheniqs version 2.1.0 [Galanti et al. 2021]. The entire process was executed using a custom nextflow pipeline, GENEFLOW [https://github.com/gencorefacility/GENEFLOW].

### Barcode verification of CNV strains and revertant clones

Of the 15 CNV strains, 12 strains (*G_4 - GM_15*) had unique barcode sequences, which we used to confirm that each strain was a unique lineage that arose independently. We screened for the barcode region using colony PCR and agarose gel electrophoresis, followed by DNA purification and quantification before Sanger sequencing. We obtained the unique barcode sequence for each strain by Clustal omega alignment on the Sanger sequences, and compared them to the known barcode template to find the varying bases.

For revertant clones, we used the barcode to confirm parental lineage and eliminate any possibility of contamination. We detected barcodes directly from whole-genome sequencing data. Across FASTQ files generated from all 56 revertants, we wrote a custom script using a global regular expression (grep) command to search for sequences matching the barcode template. The output defined the exact barcode sequence(s) present in each FASTQ file, and the number of corresponding reads. This allowed us to eliminate any barcodes present in very low frequencies, which are false positives due to index switching and not genuine detection in the genome sequence.

### Gene copy number and breakpoint analysis

For each CNV and revertant strain, we estimated gene copy number using samtools [Danecek et al. 2021] to calculate the read depth of each nucleotide from whole-genome sequencing data. We calculated read depth after aligning the reads of each strain using BWA-MEM to a reference genome that includes the reporter gene constructs present in its ancestor.

To infer potential breakpoints in CNV and revertant strains, we used our custom analysis tool CVish (https://github.com/pspealman/CVish) [Spealman et al. 2025], which takes in FASTQ files of strains of interest and generates split reads and discordant reads by comparing them to the ancestral genome. We use these to locate amplified regions in the genome and to infer CNV formation and loss mechanisms. CVish also calculates relative read depth across the genome, outputting it as a bedgraph file which shows copy number differences among different genomic regions.

### Long-read whole-genome sequencing

We extracted genomic DNA using Qiagen 20/G Genomic Tips, following the manufacturer’s protocol for yeast. We quantified gDNA concentration on the Qubit fluorometer and assessed sample quality using the Agilent Genomic Tapestation. 0.45X AMPURE PB were used for DNA cleanup and the Hologic Diagenode Megaruptor 3 for DNA shearing. DNA libraries were prepared using the PacBio SMRTbell® prep kit 3.0, according to the manufacturer’s protocol. Samples were barcoded and pooled equally. Polymerase was loaded using the PacBio Revio polymerase kit, following the manufacturer’s instructions. PacBio HiFi sequencing was done using PacBio Revio SMRT Cell. Demultiplexing was done using SMRT Link v13.1.

### Long-read data analysis and de novo genome assembly

For each sequenced strain, we converted the raw BAM files to FASTQ files using bedtools and then aligned the FASTQ files to our reference genome using the mapping argument for PacBio in minimap2 [Li 2018]. We converted the SAM output files into sorted and indexed BAM files using samtools, and visualised them on Integrated Genome Viewer (IGV) [Robinson et al. 2011]. We manually compared each revertant to its parental CNV strain at the breakpoint regions (identified by CVish from short-read data) to check for any remnant genomic trace of the CNV reflected as changes in the nucleotide sequence. We also searched for chimeric or split reads using samtools ‘view’. We used bedtools ‘genomecov’ to calculate sequencing depth for each nucleotide normalized to the genome-wide mean [Quinlan and Hall 2010], as a measure of the copy number, and checked for read peaks compared to the ancestor. Split reads extracted by samtools around identified breakpoints were assembled into contigs using miniasm (default settings, v0.3) [Li 2016; Danecek et al. 2021]. Contigs were manually verified using select reads that spanned both chromosomes.

### Single nucleotide variation analysis

For every ancestral, CNV and revertant strain, we used whole-genome sequencing data to identify genomic variations other than CNVs, including single nucleotide polymorphisms and DNA insertions and deletions. We used the bioinformatics pipeline developed by the Genomics Core at NYU CGSB (https://gencore.bio.nyu.edu/variant-calling-pipeline-gatk4/), using the Genome Analysis Toolkit 4 (GATK4) to perform variant calling and filtering. We then employed SnpEff to annotate and predict variant effects using identified SNVs, generating vcf files as the output.

GATK4 predictions typically suffer from high false-positive rates. To eliminate false-positive calls, we applied a custom pipeline (https://github.com/pspealman/heranca) [Spealman et al. 2025], that flags likely low-quality predictions arising from homopolymer or polynucleotide runs and identifies mutations that break the infinite sites assumption by occurring in at least three unrelated strains as likely false positives.

As a final step to get high-confidence SNVs, we individually evaluated SNVs by visualizing the bam files on Integrated Genome Viewer (IGV) [Robinson et al. 2011]. We compared each SNV in a revertant strain to the same position on its CNV parent, as well as its one-copy ancestor strain. Only SNVs exclusively present in one strain, with a base conversion frequency of 90% or higher of total reads, were designated as high-confidence SNPs.

### Evolutionary model and simulation-based inference

We modeled the experiments using a Wright-Fisher model with selection, mutation, and drift. First, we considered a scenario without epistasis between *GAP1* and *MEP2* CNV reversions. Without epistasis, we can consider each CNV reversion as an independent allele, with its own selection coefficient *s* (*s_G_ or s_M_*), reversion rate δ (δ*_G_ or* δ*_M_*), and initial frequency φ (φ*_G_ or* φ*_M_*). Then, we infer the parameters of each experimental replicate separately. Thus, we infer three parameters per mutation, for a total of six parameters per replicate. We assume CNV reversion to be either beneficial or effectively neutral, as the fitness of CNV reversion is 1 + *s* > 1. Then, estimating the fitness effect of CNVs, *w*_*cnv*_, given *s* is 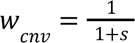. The overall fitness of a strain was calculated assuming multiplicative fitness., 1 + *s_GM_* = (1 + *s_G_*)×(1 + *s_M_*). We used Neural Posterior Estimation (NPE; Greenberg et. al 2019, Papamakarios and Murray 2016) with Masked Autoregressive Flow (Papamakarios et. al 2017) for parameter inference, as in our previous studies [Avecilla et al 2022, Chuong et al 2025]. The training set consists of 10,000 evolutionary simulations and their parameter values, and the neural network is trained using a log-likelihood loss to approximate the amortized posterior distribution.

To bridge the gap between synthetic simulations and empirical observations we added a small Gaussian noise to each simulation, such that the simulation result given a parameter set θ, *x*(θ), becomes *x*_*noi*_*_s_*_*y*_(θ) = *x*(θ) + ε; ε∼*N*(0, 0. 02). Note that for dynamics without epistasis we only require one trained neural network that can estimate the posterior distributions of both *GAP1* and *MEP2* CNV reversion parameters. We used a log-uniform prior for parameter values (**Supplemental Table 4**). Then, we computed the collective posterior distribution, a single posterior distribution conditioned on all replicates of each strain [see Chuong et al. 2025 for full details]. To validate the estimation of the individual and collective posterior distributions, we performed posterior predictive checks that measure the similarity between simulations using posterior parameter samples and the empirical observations.

## Data availability

Sequencing data is available at SRA PRJNA1291234. The source code repository for simulation-based inference is on Github: https://github.com/nadavbennun1/de_et_al. Scripts for all other analysis and figures, as well as all data, are also available on Github: https://github.com/GreshamLab/CNV_Stability_without_selection_pressure.

## Acknowledgements

We thank members of the Gresham and Siegal labs for discussions and feedback. Funding for this research was provided by NIGMS (R35GM153419) and the US - Israel Binational Science Foundation (2021276). This work was supported in part through the NYU IT High Performance Computing resources, services, and staff expertise. We acknowledge the Zegar Family Foundation for their generous support. We thank the NYU Center for Genomics and System Biology Genomics Core for their assistance and resources.

## Supplementary Figures

**Figure S1.**
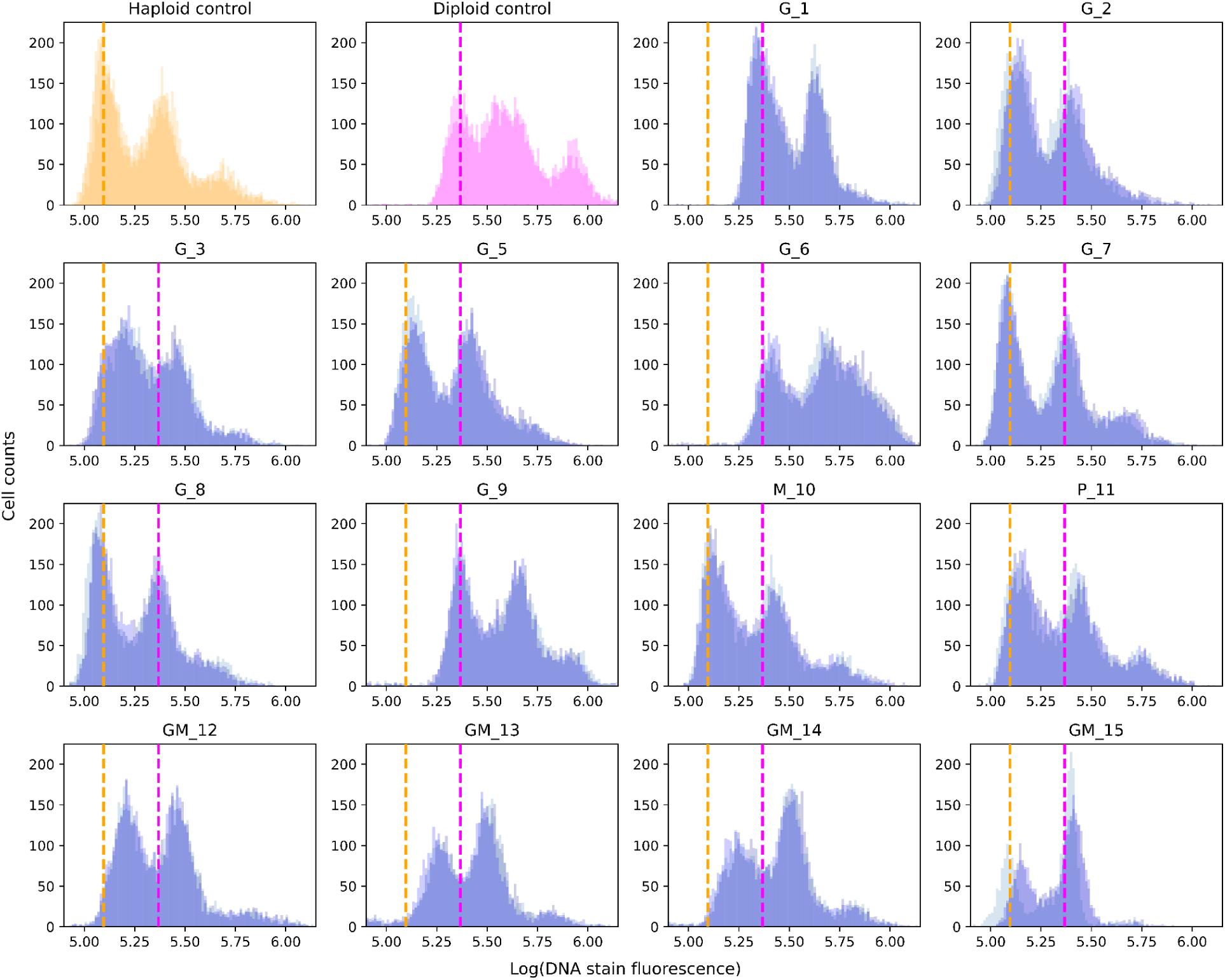
Ploidy of all CNV strains used in the study. Ploidy was measured by quantifying DNA content using propidium iodide staining of exponentially growing cells. Haploid (orange) and diploid (magenta) control strains were assayed to define ploidy states and all strains were tested in triplicate (overlaid in each subplot). Dashed lines indicate the modes calculated from all replicates of each control, and mark the position of the 1N peak for haploids (orange) and the 2N peak for diploids (magenta). All CNV strains are shown here except *G_4* (shown in Figure S2).

**Figure S2.**
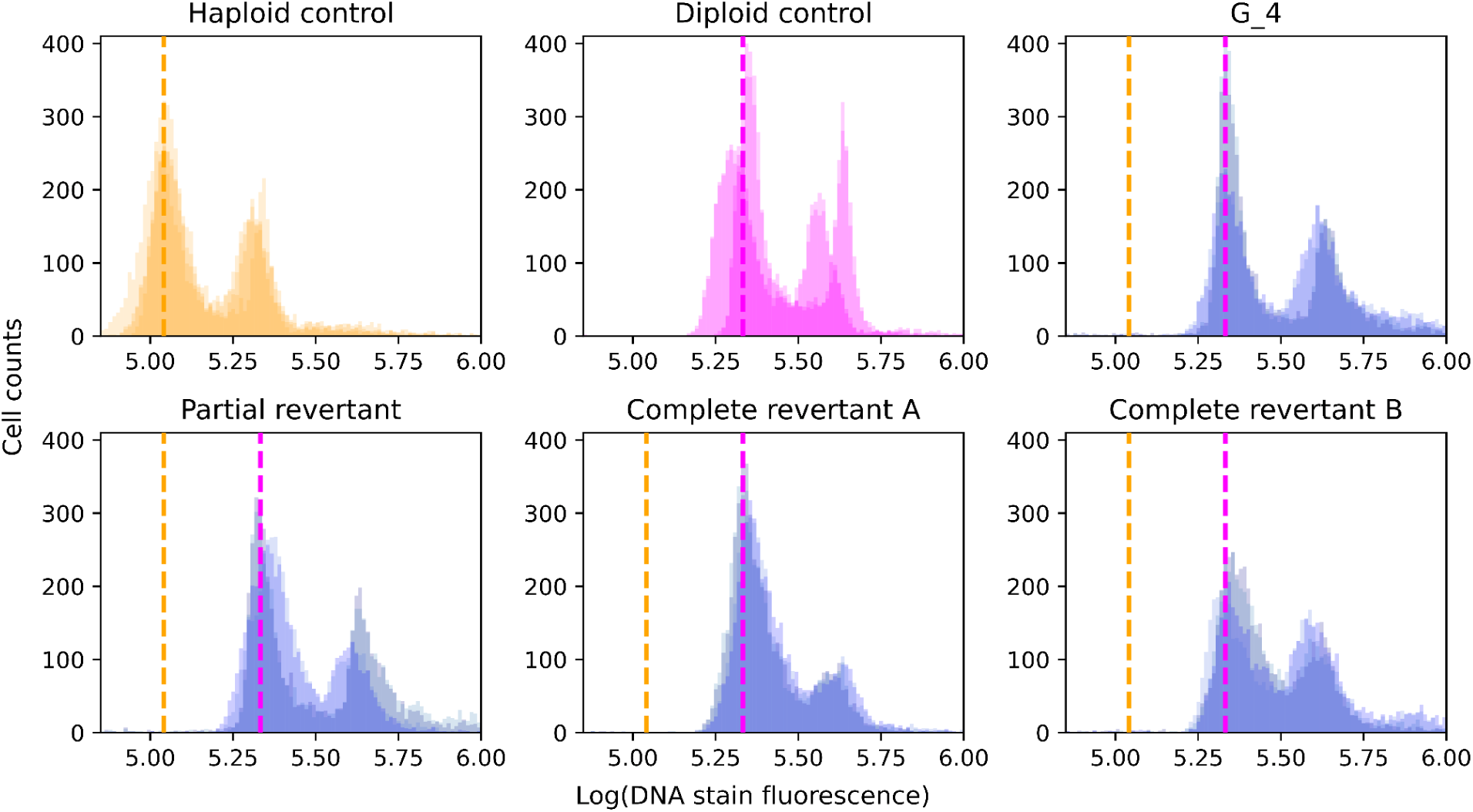
Ploidy of segmental CNV strain *G_4* and derived revertants. Ploidy was measured by quantifying DNA content using propidium iodide staining of exponentially growing cells. Haploid (orange) and diploid (magenta) control strains were assayed to define ploidy states and all strains were tested in quadruplicate (overlaid in each subplot). Dashed lines indicate the modes calculated from all replicates of each control, and mark the position of the 1N peak for haploids (orange) and the 2N peak for diploids (magenta).

**Figure S3.**
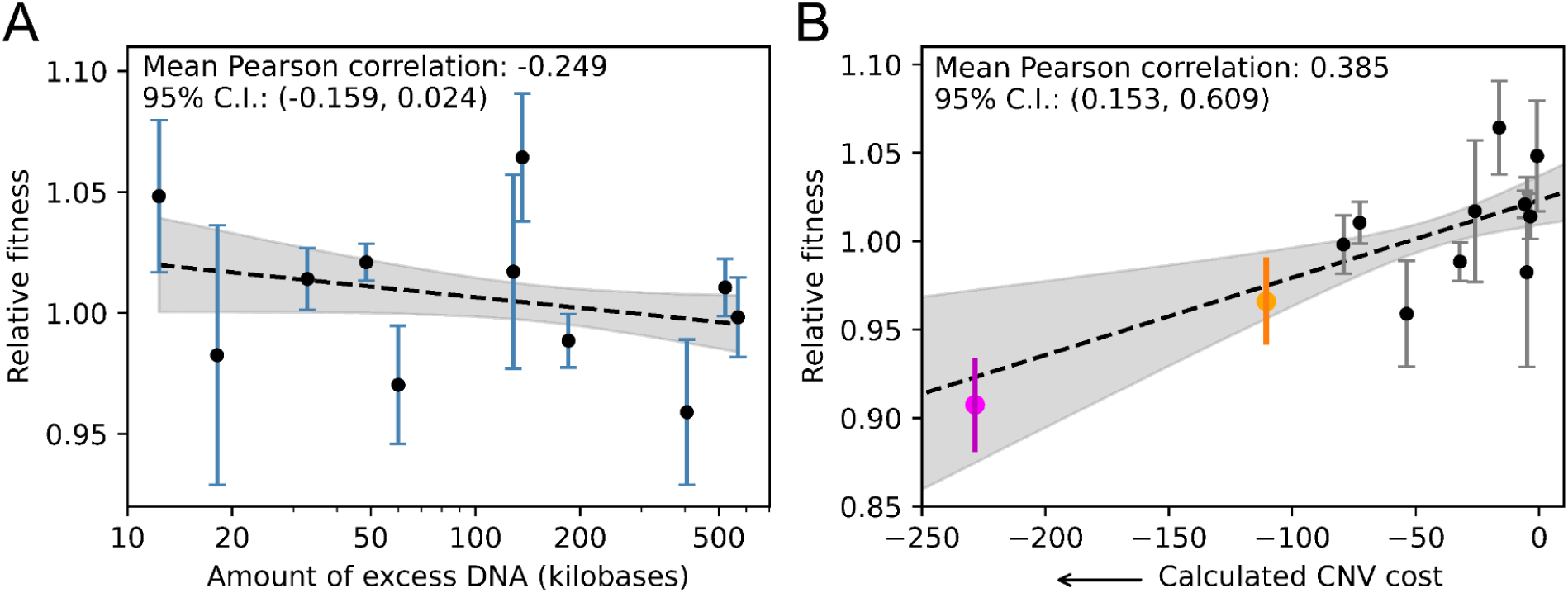
Relationship of relative fitness with amount of excess DNA and with estimated CNV cost. **A)** Pearson correlation coefficient was calculated between relative fitness and amount of extra DNA in all segmental CNVs. Points with error bars indicate 95% confidence intervals for relative fitness. Black dashed line and grey shadow indicate 95% confidence intervals for a linear regression through all points. Relative fitness has no significant correlation (95% CI: −0.159, 0.024) with the amount of excess DNA. **B)** Relative fitness has significant correlation (95% CI: 0.153, 0.609) with estimated fitness costs of segmental CNVs. Double aneuploidy of chromosomes XI and XIV (magenta) and single aneuploidy of chromosome XI (orange) lie on the regression line fitted to all segmental CNVs.

**Figure S4.**
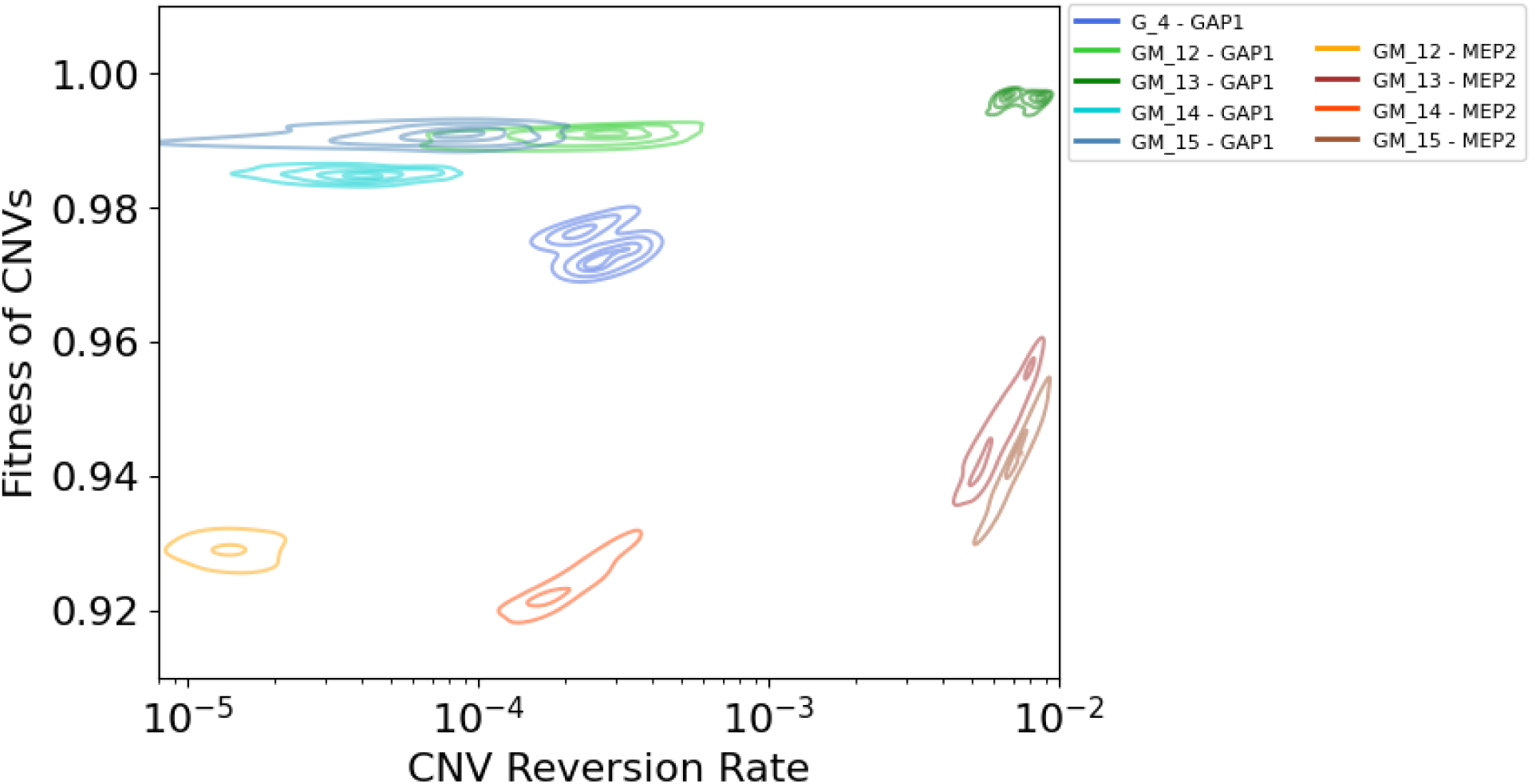
Parameter estimates. 95% high-density regions (HDRs) of the collective posterior distributions for the fitness cost (derived from selection coefficients, see Methods) and reversion rate of CNVs.

**Figure S5.**
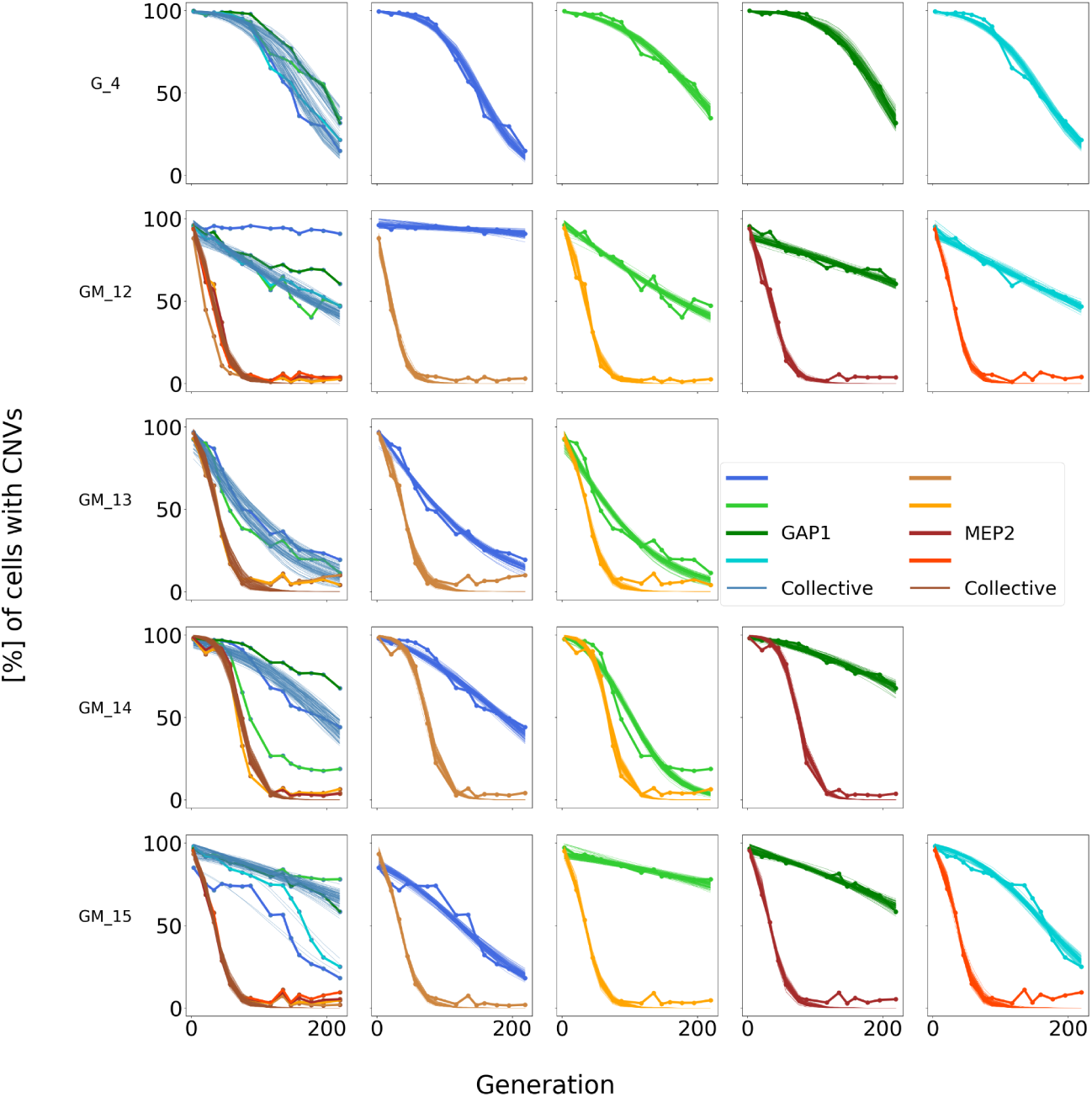
Posterior predictive checks. Model simulations of 100 parameter sets sampled from the posterior distributions inferred from each experimental replicate (shaded lines) against the empirical data (solid lines with markers). Leftmost (grey) panels of each row show predictive checks of the collective posterior distribution for the strain, i.e., a posterior distribution conditioned on all strain replicates.

**Figure S6.**
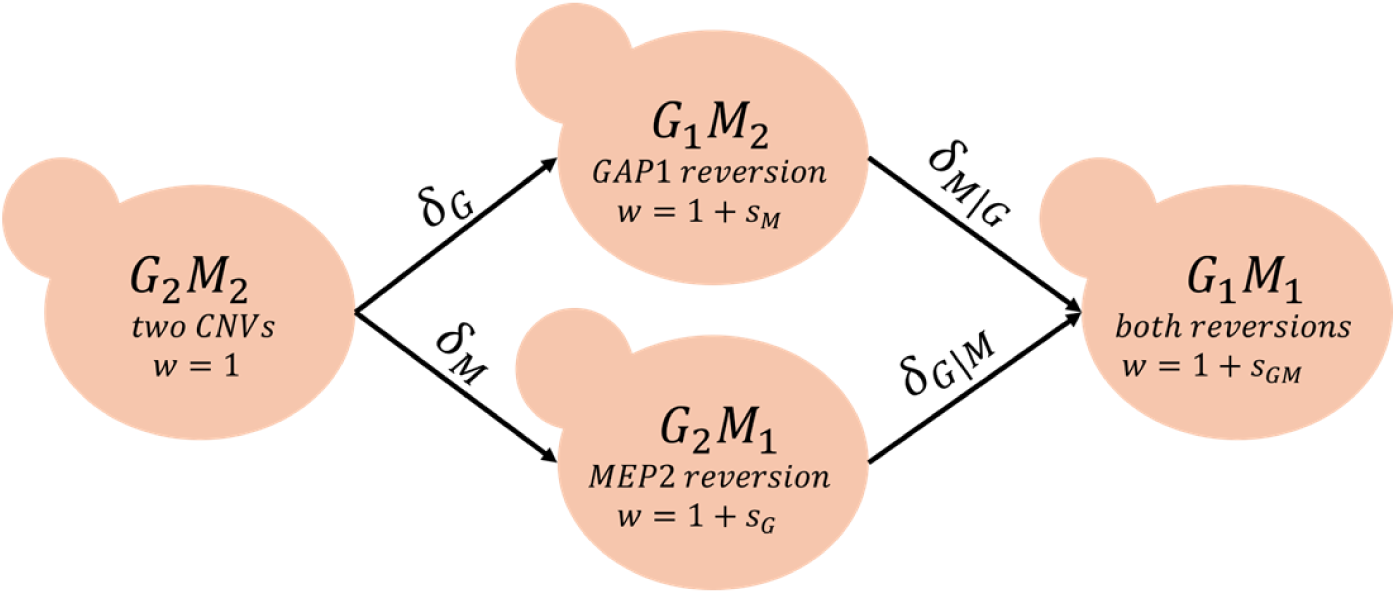
A genotype-tracking evolutionary model. CNV reversions are not necessarily independent, with rates δ_*i*|*j*_ for transitioning from genotype *j* to genotype *i*, and a distinct selection coefficient *s*_*k*_ for genotype *k*. We track the proportions of each genotype, from which we can calculate the total proportions of each CNV. Cell labels refer to both genes (G - *GAP1*, M - *MEP2*) and their corresponding copy numbers, i.e., indicate the entire cell’s genotype.

**Figure S7.**
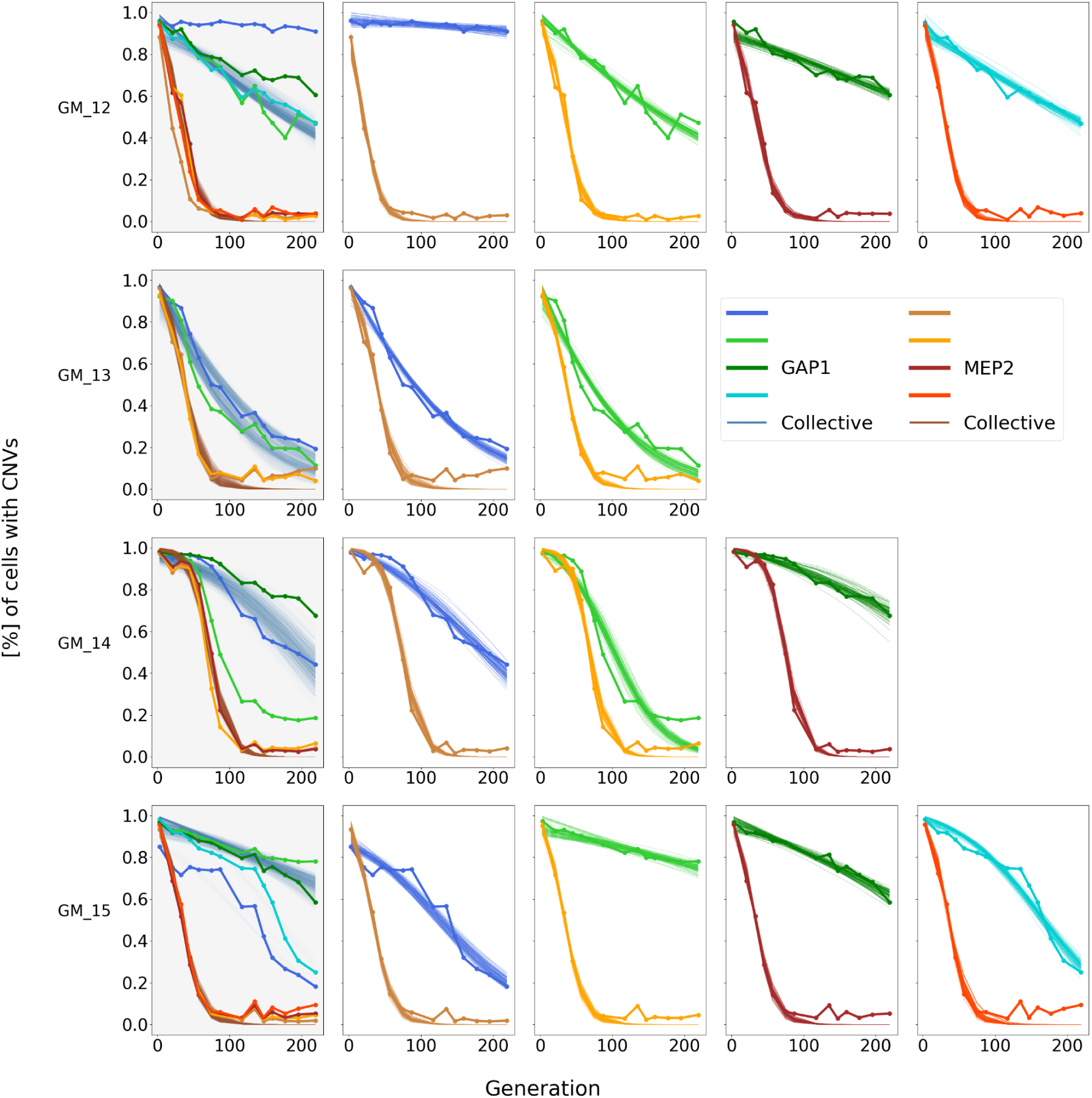
Posterior predictive checks with the genotype-tracking model. Genotype-tracking model simulations of 100 parameter sets sampled from the posterior distributions of each experimental replicate (shaded lines), inferred using the allele-tracking model against the empirical data (solid lines with markers). Leftmost (grey) panels of each row show predictive checks of the collective posterior distribution for the strain, i.e., a posterior distribution conditioned on all strain replicates. Strain *G_4* is not shown since it does not have a *MEP2* CNV.

**Figure S8.**
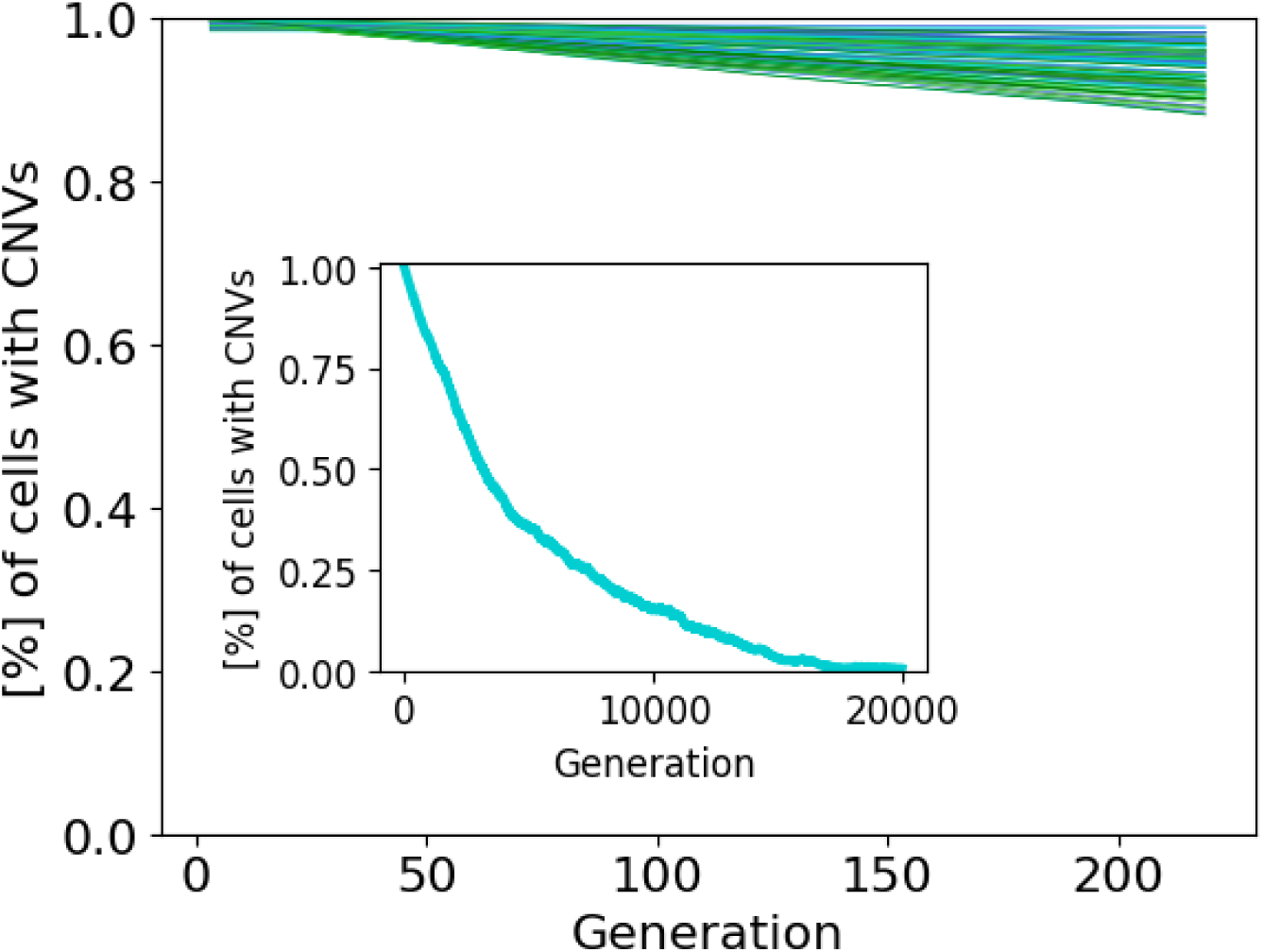
*GAP1* CNV reversion without selection. Evolutionary simulations with mutation rate and initial CNV proportion sampled from the collective posterior distribution of strain *G_4* and with very weak selection, *s* = 10^−10^. Inset axis shows a simulation of the sample mean for 20,000 generations.

**Figure S9.**
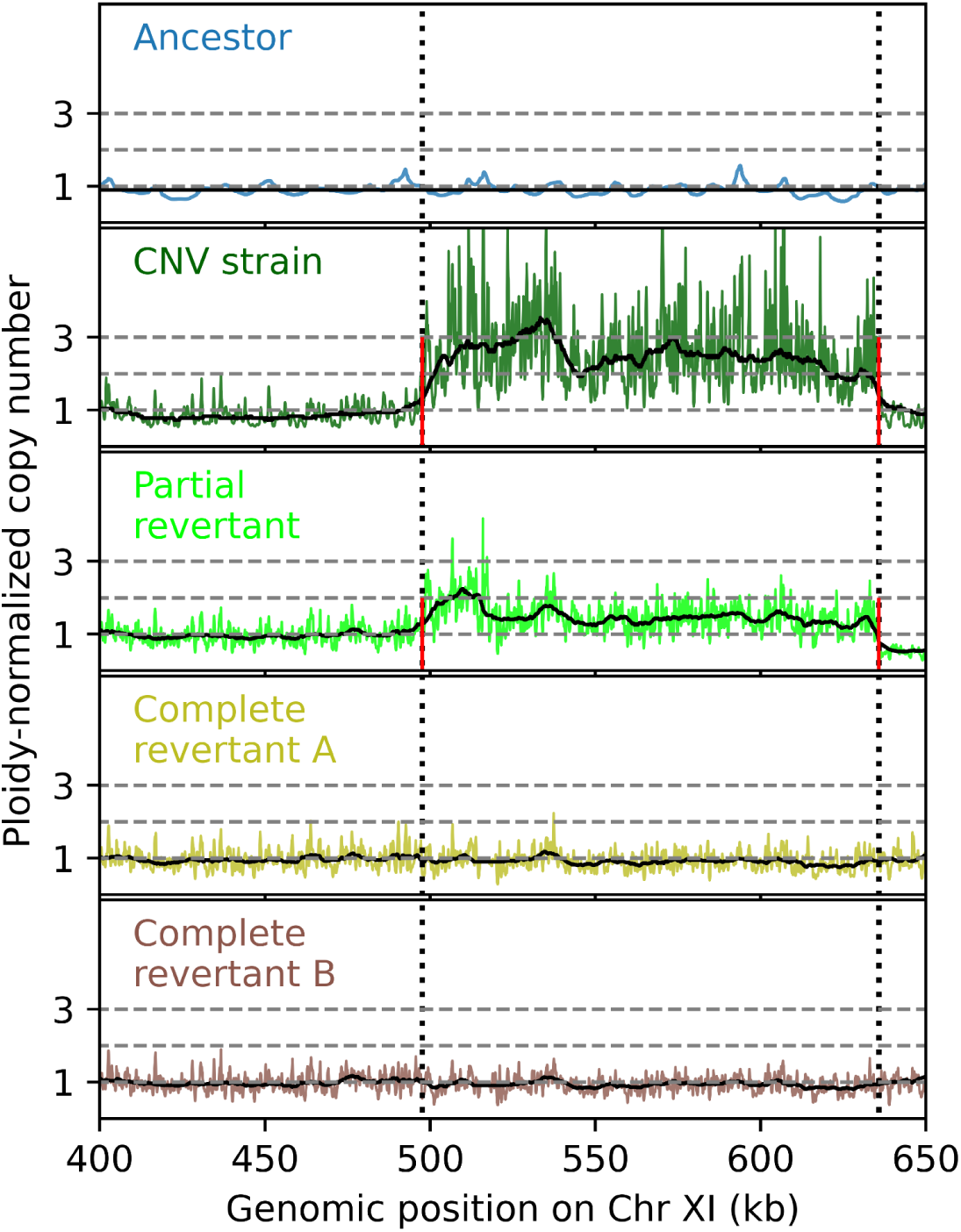
Short-read whole-genome sequencing of segmental CNV strain *G_4* and derived revertants. Sequencing read depth for each nucleotide normalized to the genome-wide mean, and averaged over all copies of the chromosome (1 for haploids, 2 for diploids), is the ploidy-normalized copy number (PNCN). Split reads (in red) indicate CNV breakpoints. CNV boundaries are marked by vertical dotted black lines; horizontal dashed grey lines indicate PNCN of 1, 2 and 3 for visual comparison. Solid black lines indicate median read depth over rolling 50 kb windows. All strains except the ancestor are diploid. We compared the CNV strain *G_4* to its ancestor, a partial revertant, and 14 complete revertants (two are shown here).

**Figure S10.**
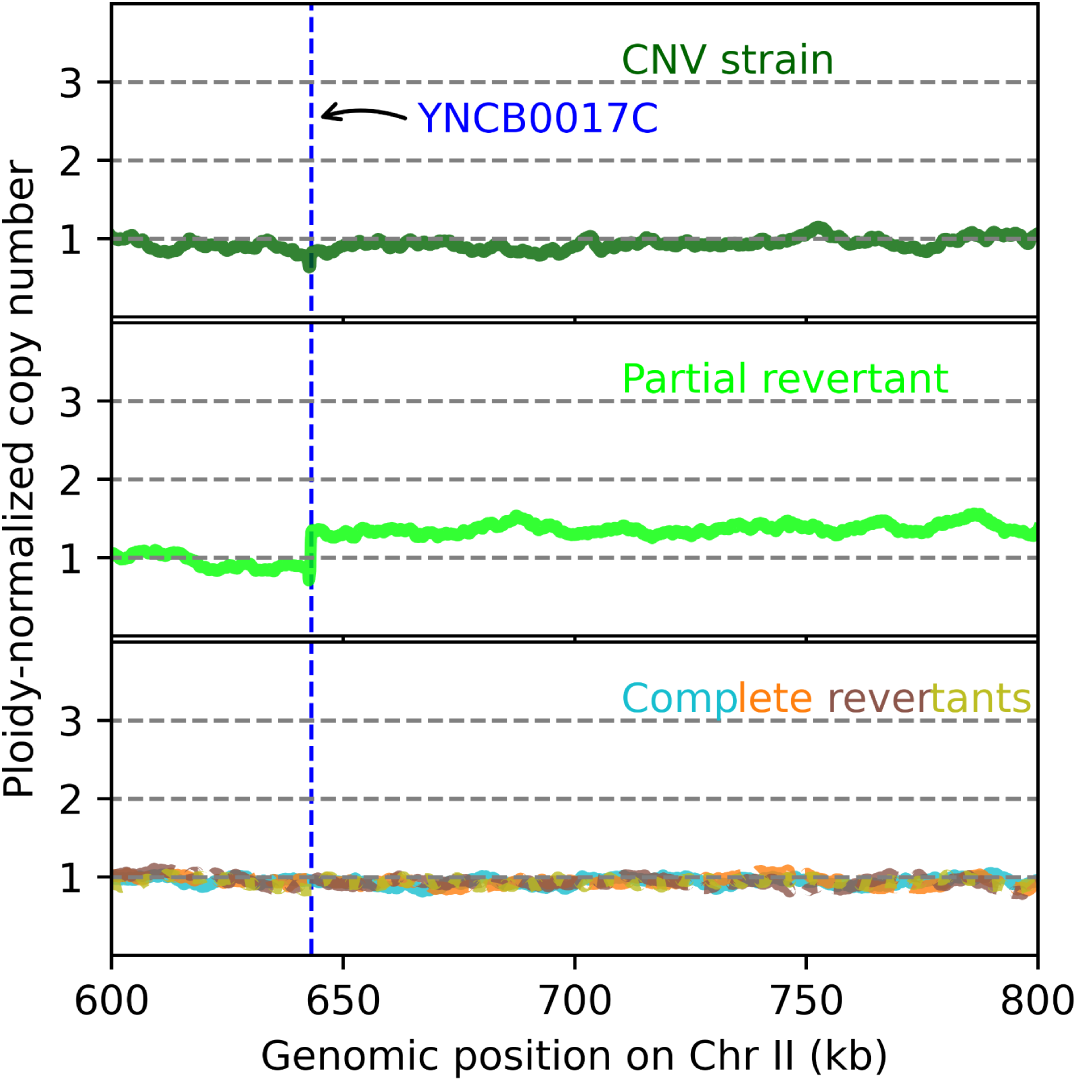
Long-read sequencing showing ploidy-normalized copy number across a part of chromosome II for the segmental CNV strain *G_4* and derived revertants. Sequencing read depth for each nucleotide normalized to the genome-wide average, and averaged over two copies of the chromosome for diploids, is the ploidy-normalized copy number (PNCN). Horizontal dashed grey lines indicate PNCN of 1, 2 and 3 for visual comparison. The vertical dashed blue line indicates the position of the *YNCB0017C* gene located at the boundary of the region translocated to the neochromosome. Shown here are the segmental CNV that underwent reversion (*G_4*), a partial revertant, and four independently evolved complete revertants (overlaid) – all diploids.

**Figure S11.**
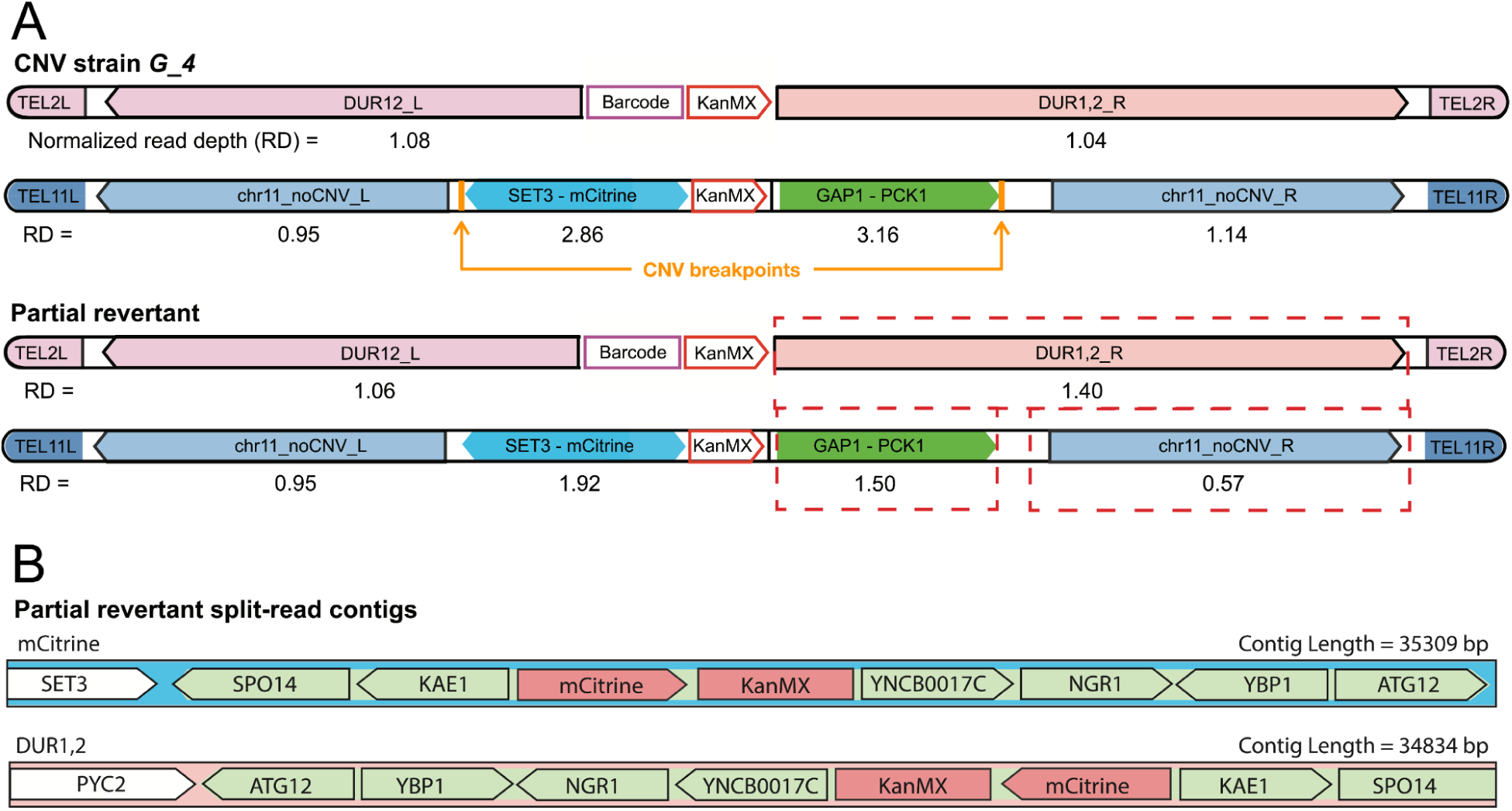
De novo genome assembly and ploidy-normalized copy number reveal mechanism of partial reversion. **A)** Sequencing read-depth of different parts of the amplified region relative to the genome-wide average, and then normalized by ploidy. **B)** Two contigs obtained by de novo genome assembly, which span both chromosomes XI and II.

**Figure S12.**
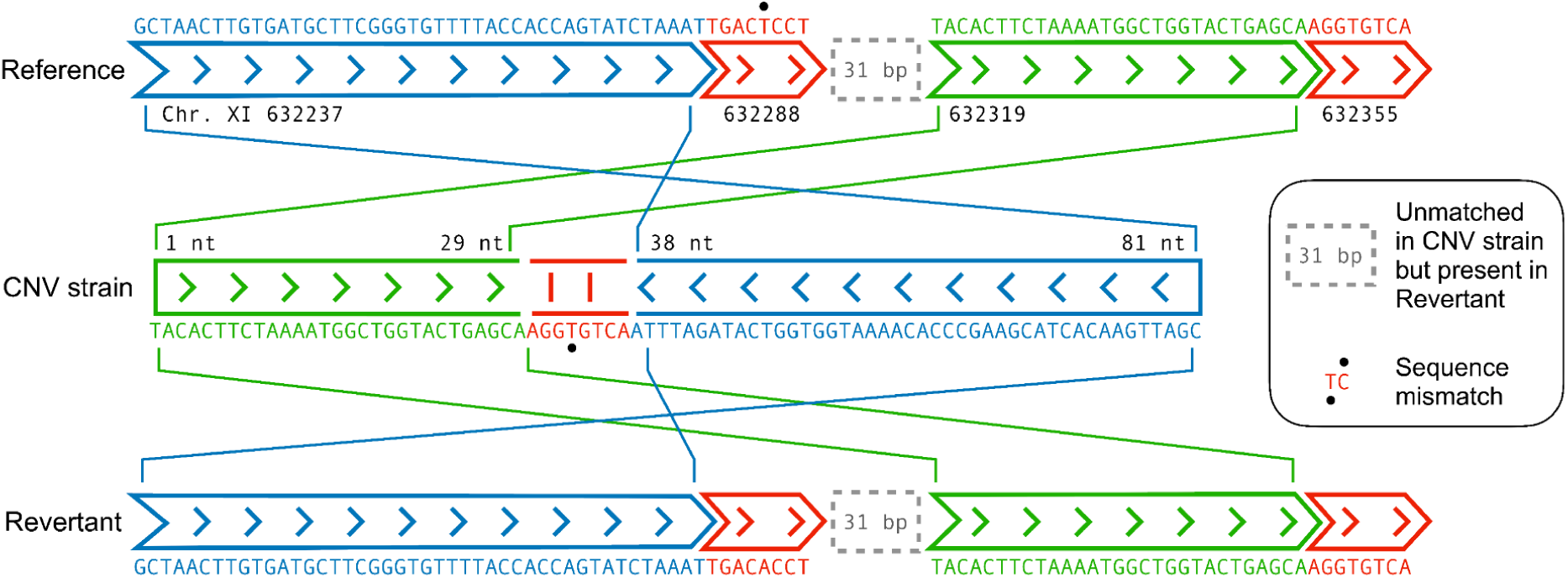
The segmental amplification, bearing an inversion in the parent CNV strain *G_4*, returns to the identical ancestral state in all 14 complete revertants (right-hand breakpoint of one example is shown here).

## Supplementary Tables

**Table 1.**
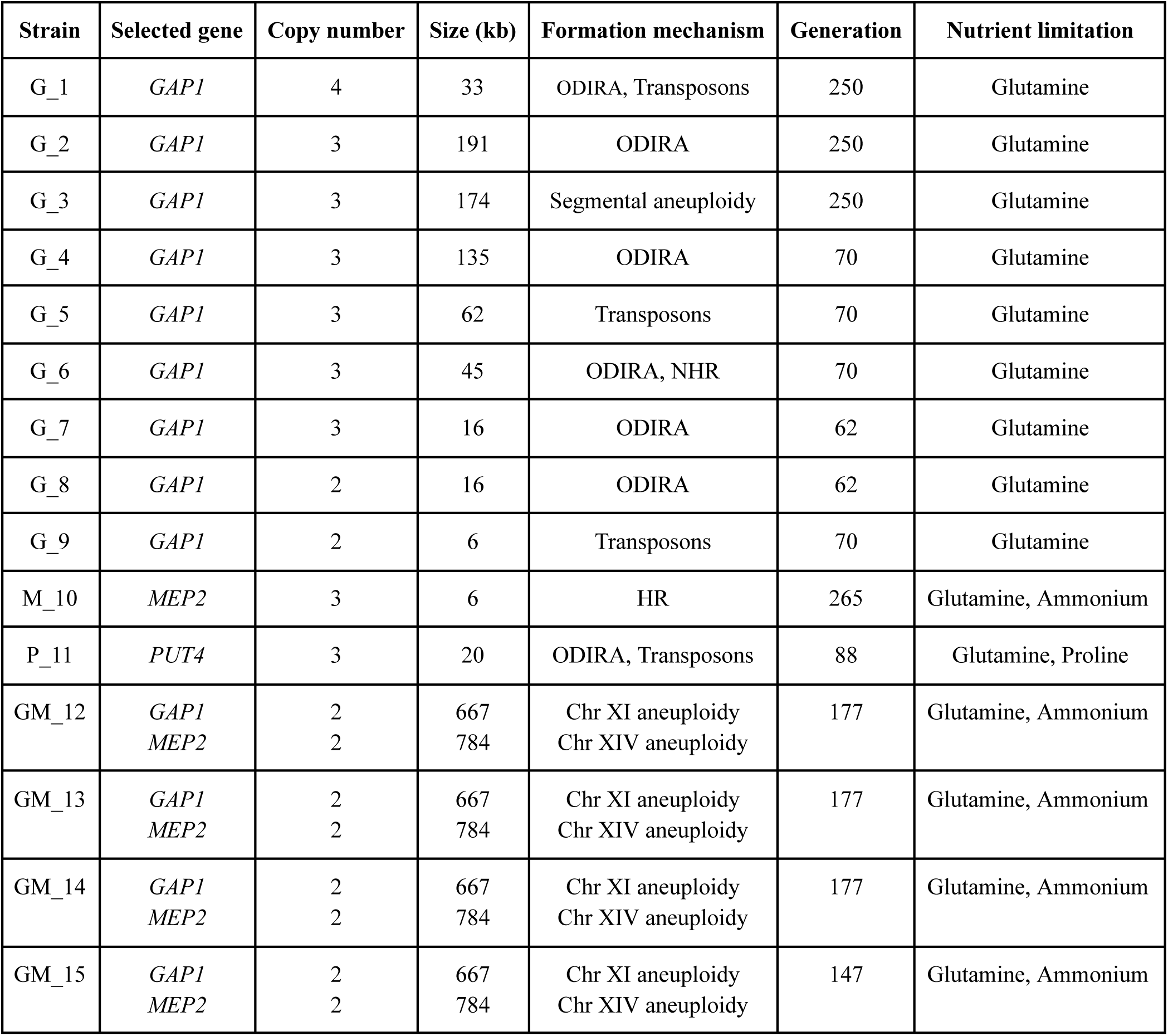
CNV strains used in this study. Summarized here are the different parameters by which the CNVs differed: the gene(s) under selection, copy number relative to the ancestor, size of the amplified region, molecular mechanism underlying CNV formation, number of elapsed generations before the lineage was isolated and the selection condition under which they arose.

**Table 2.** CNV strain nomenclature and origin. Lab collection ID, and publications which reported the experiments from which each strain originated.

| **Name of strain in the paper** | **Lab collection ID** | **Strain origin** |
| --- | --- | --- |
| *G_1* | DGY1741 | Lauer et al. (2018) |
| *G_2* | DGY1743 | Lauer et al. (2018) |
| *G_3* | DGY1744 | Lauer et al. (2018) |
| *G_4* | DGY1886 | Lauer et al. (2018) |
| *G_5* | DGY1917 | Lauer et al. (2018) |
| *G_6* | DGY1883 | Lauer et al. (2018) |
| *G_7* | DGY1988 | Lauer et al. (2018) |
| *G_8* | DGY1976 | Lauer et al. (2018) |
| *G_9* | DGY1920 | Lauer et al. (2018) |
| *M_10* | DGY2702 | Abdul-Rahman et al. (in prep) |
| *P_11* | DGY2769 | Abdul-Rahman et al. (in prep) |
| *GM_12* | DGY2748 | Abdul-Rahman et al. (in prep) |
| *GM_13* | DGY2750 | Abdul-Rahman et al. (in prep) |
| *GM_14* | DGY2751 | Abdul-Rahman et al. (in prep) |
| *GM_15* | DGY2755 | Abdul-Rahman et al. (in prep) |

**Table 3.**
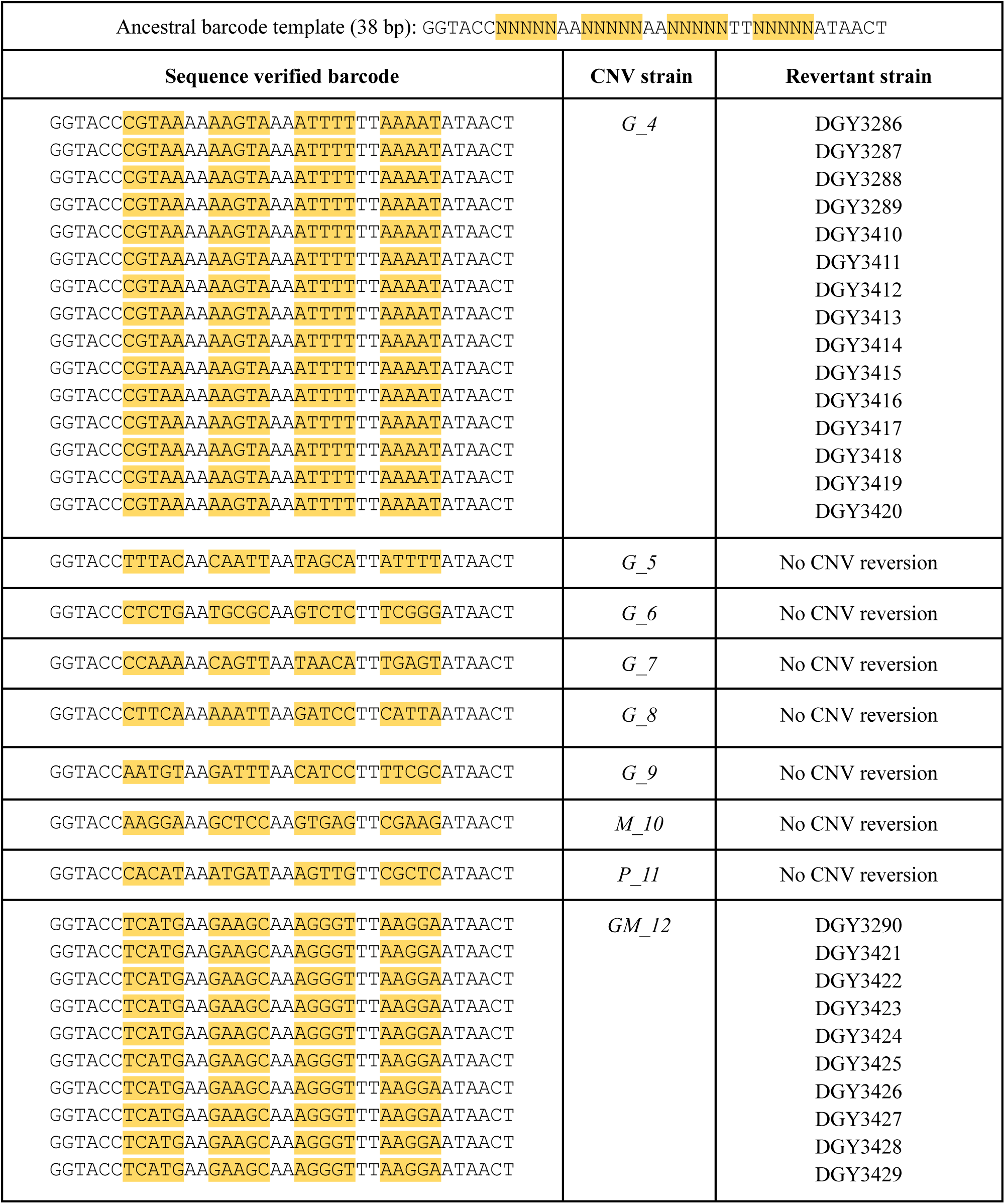
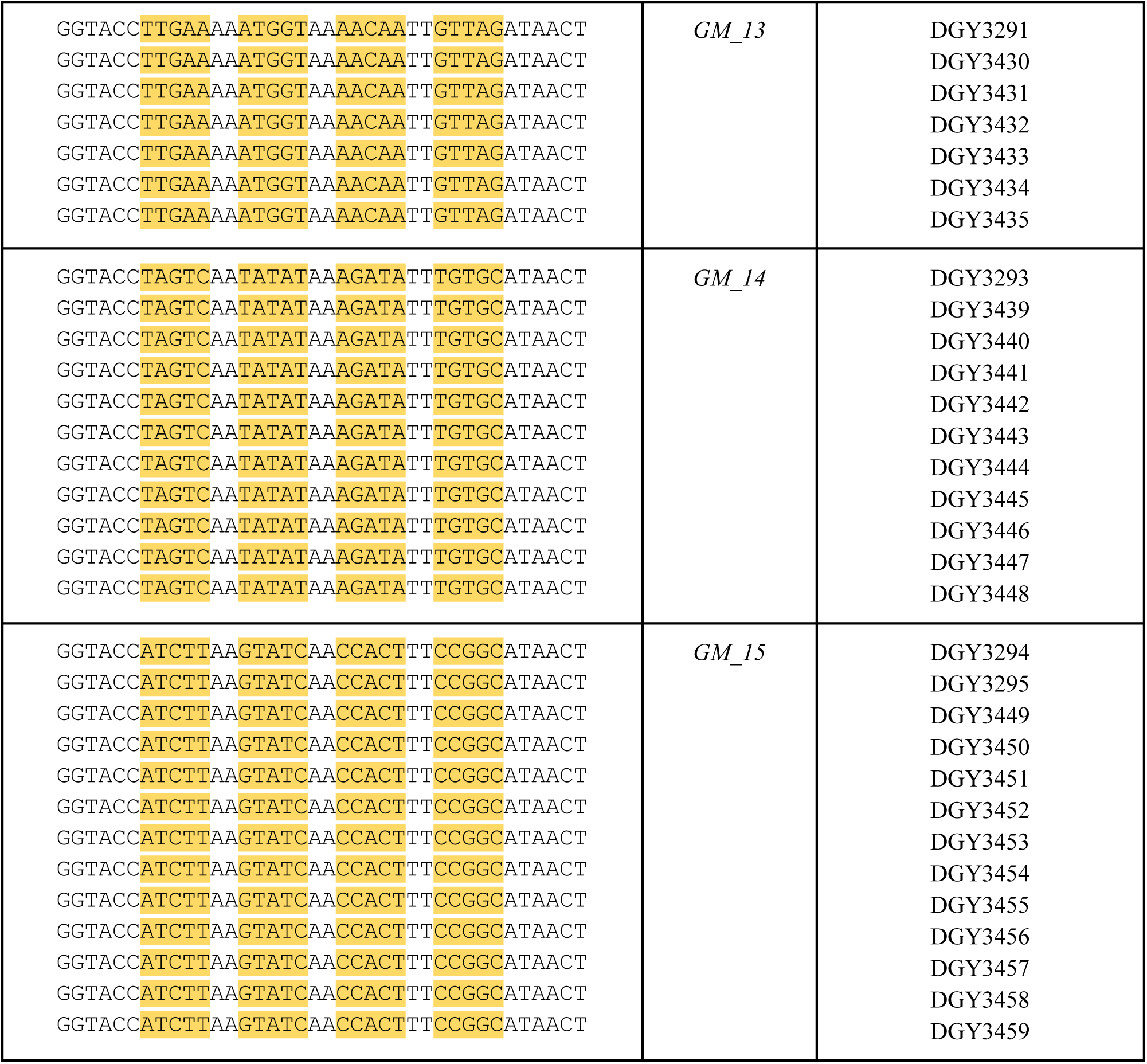
Unique barcodes identifying each CNV lineage and revertants descended from them. Each sequence was obtained from whole genome sequencing data by searching for the ancestral barcode template. Variable sequence regions in each barcode are highlighted.

**Table 4.** Marginal prior distributions of the evolutionary parameters. The overall prior distribution is the hypercube defined by these uniform marginal distributions.

| Parameter | Description | Prior |
| --- | --- | --- |
| $s$ | Selection coefficients of CNV reversion | $\log_{10}(s) \sim U[-3, -1]$ |
| $\delta$ | CNV reversion rate | $\log_{10}(\delta) \sim U[-5, -2]$ |
| $\varphi$ | Initial frequency of reversed CNV cells | $\log_{10}(\varphi) \sim U[-5, -0.5]$ |

**Table 5.** Confirmed SNVs across all CNV strains and revertants. For each SNV, we record the genomic position, the original base and the new base to which it changed, the percentage of reads which underwent the base change, the gene or intergenic region where the SNV is located, and whether the revertant underwent partial (lost only Chr XIV) or complete (lost both Chr XI and Chr XIV) reversion. We did not detect any high-confidence SNVs in any of the 15 CNV strains used in the study.

| CNV Strain | Revertant strain | Chr | Position (bp) | Original base | New base | Percentage of reads | Gene or intergenic region | Reversion type |
| --- | --- | --- | --- | --- | --- | --- | --- | --- |
| GM_12 | GM_12_Rev1 | VII | 245951 | A | G | 94% of 140 | FLC3 | Complete |
|  | GM_12_Rev1 | XIII | 712911 | C | T | 95% of 169 | ERG8 | Complete |
|  | GM_12_Rev2 | I | 54719 | G | A | 95% of 138 | GEM1 | Complete |
|  | GM_12_Rev6 | IV | 675218 | C | T | 94% of 63 | YDR109C | Complete |
|  | GM_12_Rev10 | XV | 626108 | C | A | 96% of 77 | SLP1 | Partial |
| GM_13 | GM_13_Rev1 | X | 536002 | C | G | 96% of 56 | Between BFA1 and KCH1 | Complete |
|  | GM_13_Rev2 | XI | 669969 | G | A | 92% of 187 | Near GEX2 | Complete |
|  | GM_13_Rev6 | V | 530182 | A | T | 96% of 117 | RAD3 | Complete |
| GM_14 | GM_14_Rev1 | XIII | 160549 | G | A | 94% of 101 | CMP2 | Complete |
|  | GM_14_Rev1 | XIII | 160563 | A | G | 97% of 106 | CMP2 | Complete |
|  | GM_14_Rev2 | III | 226561 | T | C | 94% of 89 | TVS1 (YCR061W) | Complete |
|  | GM_14_Rev10 | XVI | 604956 | C | A | 100% of 96 | SDD4 (YPR022C) | Partial |
| GM_15 | GM_15_Rev1 | X | 517755 | A | T | 98% of 127 | Between POL32 and IMT3 | Complete |
|  | GM_15_Rev3 | VIII | 170137 | G | A | 97% of 153 | SLT2 | Complete |
|  | GM_15_Rev6 | IV | 1171363 | C | T | 96% of 142 | PAL1 | Partial |
|  | GM_15_Rev6 | V | 490478 | C | T | 93% of 184 | Between tE(UUC)E3 and YER158C | Partial |
|  | GM_15_Rev7 | XI | 71825 | A | T | 93% of 117 | PEX1 | Complete |
|  | GM_15_Rev8 | VIII | 240308 | T | C | 98% of 135 | ERG7 | Complete |
|  | GM_15_Rev8 | XV | 410348 | C | T | 95% of 105 | Between snR62 and WHI2 | Complete |
|  | GM_15_Rev12 | IV | 645268 | C | G | 95% of 129 | Between tQ(UUG)D2 and YDR098C-B | Complete |

**Table 6.** Strain nomenclature for all revertants. Names of all revertant as used in the paper, and their corresponding lab collection ID.

| **Revertant** | **Lab collection ID** |  | **Revertant** | **Lab collection ID** |
| --- | --- | --- | --- | --- |
| G_4_Rev1 | DGY3286 |  | GM_13_Rev4 | DGY3432 |
| G_4_Rev2 | DGY3287 |  | GM_13_Rev5 | DGY3433 |
| G_4_Rev3 | DGY3288 |  | GM_13_Rev6 | DGY3434 |
| G_4_Rev4 | DGY3289 |  | GM_13_Rev7 | DGY3435 |
| G_4_Rev5 | DGY3410 |  | GM_14_Rev1 | DGY3293 |
| G_4_Rev6 | DGY3411 |  | GM_14_Rev2 | DGY3439 |
| G_4_Rev7 | DGY3412 |  | GM_14_Rev3 | DGY3440 |
| G_4_Rev8 | DGY3413 |  | GM_14_Rev4 | DGY3441 |
| G_4_Rev9 | DGY3414 |  | GM_14_Rev5 | DGY3442 |
| G_4_Rev10 | DGY3415 |  | GM_14_Rev6 | DGY3443 |
| G_4_Rev11 | DGY3416 |  | GM_14_Rev7 | DGY3444 |
| G_4_Rev12 | DGY3417 |  | GM_14_Rev8 | DGY3445 |
| G_4_Rev13 | DGY3418 |  | GM_14_Rev9 | DGY3446 |
| G_4_Rev14 | DGY3419 |  | GM_14_Rev10 | DGY3447 |
| G_4_Rev15 | DGY3420 |  | GM_14_Rev11 | DGY3448 |
| GM_12_Rev1 | DGY3290 |  | GM_15_Rev1 | DGY3294 |
| GM_12_Rev2 | DGY3421 |  | GM_15_Rev2 | DGY3295 |
| GM_12_Rev3 | DGY3422 |  | GM_15_Rev3 | DGY3449 |
| GM_12_Rev4 | DGY3423 |  | GM_15_Rev4 | DGY3450 |
| GM_12_Rev5 | DGY3424 |  | GM_15_Rev5 | DGY3451 |
| GM_12_Rev6 | DGY3425 |  | GM_15_Rev6 | DGY3452 |
| GM_12_Rev7 | DGY3426 |  | GM_15_Rev7 | DGY3453 |
| GM_12_Rev8 | DGY3427 |  | GM_15_Rev8 | DGY3454 |
| GM_12_Rev9 | DGY3428 |  | GM_15_Rev9 | DGY3455 |
| GM_12_Rev10 | DGY3429 |  | GM_15_Rev10 | DGY3456 |
| GM_13_Rev1 | DGY3291 |  | GM_15_Rev11 | DGY3457 |
| GM_13_Rev2 | DGY3430 |  | GM_15_Rev12 | DGY3458 |
| GM_13_Rev3 | DGY3431 |  | GM_15_Rev13 | DGY3459 |

## Supplementary Methods

### Evolutionary model and simulation-based inference

#### Evolutionary model (without epistasis)

The following presents a model for the evolution of *GAP1* CNV reversion. We follow the frequencies of two genotypes over time: 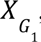, the frequency of genotype *G*_1_, representing single-copy *GAP1*; and 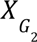, the frequency of genotype *G*_2_, representing two copies of *GAP1*.

The model for *MEP2* CNV reversion is similar, with *M* (for *MEP2*) replacing *G* (for *GAP1*).

Initially, the genotype frequencies are:

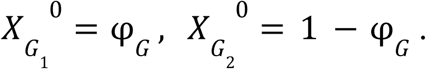

where φ*_G_* is the initial frequency of genotype *G*_1_ and is a model parameter.

The change in frequencies due to mutation is given by

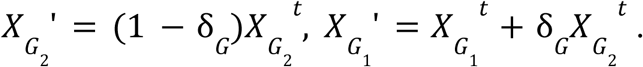

where *Xg* ’ is the frequency of genotype *g* after mutation.

The change in frequencies due to selection is given by

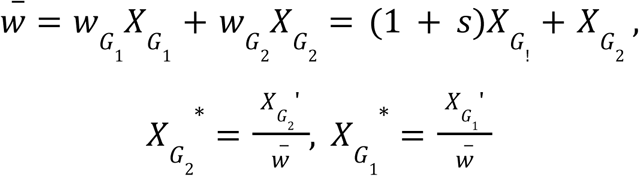

where *w*_*g*_ is the fitness of genotype *g*, and *w̄* is the population mean fitness. *X_g_*\* represents the frequency of genotype *g* after selection.

The change in frequencies due to random genetic drift is given by

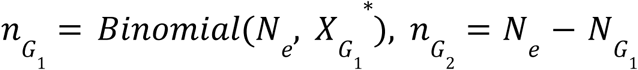

where *n*_*g*_ is the number of cells with genotype *g*.

The frequencies in the next generation are then given by

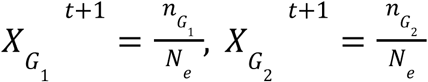

Overall, we estimate three model parameters: *s_G_*, δ*_G_*, φ*_G_*. The effective population size is constant at *N_e_* = 1.9 × 10^5^ (see details below).

The evolutionary model for *MEP2* CNV reversion is similar, with *s_M_*, δ*_M_*, φ*_M_* substituting *s_G_*, δ*_G_*, φ*_G_*, and 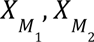 substituting 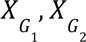

#### Effective population size

Populations begin each full growth cycle at *N*_0_ = 6. 25 × 10 cells and duplicate six times before back-dilution to *N*_0_. Using the harmonic mean [Gillespie 2004] for calculating the effective population size, we have

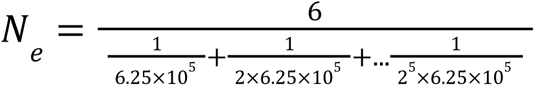

The denominator can be calculated using a geometric sum, yielding

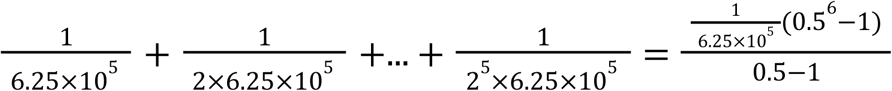

Leading to

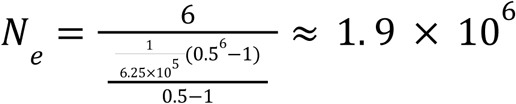

#### Evolutionary model with epistasis

The following describes a model for evolution of *GAP1* and *MEP2* CNV reversions with epistasis between the two CNVs (Figure S8_3). In this model, we consider four genotypes: *G*_2_ *M*_2_, cells with both *GAP1* CNV and *MEP2* CNV (i.e., at least one *MEP2* duplication); *G*_1_ *M*_2_, cells with a single copy of *GAP1* and a *MEP2* CNV; *G*_2_ *M*_1_, cells with a *GAP1* CNV and two copies of MEP2; and *G*_1_ *M*_1_, cells with single copies of both *GAP1* and *MEP2*. The frequency of genotype *g* at time *t* is denoted by *X_g_* ^*t*^.

Each genotype has a distinct fitness, *w_g_* = 1 + *s_g_*, with the fitness of the double CNV genotype *G*_2_ *M*_2_ fixed as 1 without loss of generality. Each of the three mutant genotypes starts with a distinct initial frequency, denoted by φ*_g_*; and the reversion rate of CNV *c* is denoted by δ*_c_*, with *c* = *M* for *MEP2* and *c* = *G* for *GAP1*. The rate of a second reversion *j* is distinct and dependent on the first reversion *i*, denoted by δ_*j*|*i*_. This model therefore has ten parameters: 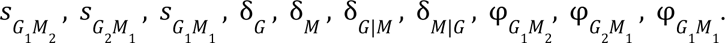

Assuming that CNV reversion rates are independent of the genotype, and that there is no epistasis between CNV reversions, we can calculate the values of the four parameters that are unique to the model with epistasis from the six parameters of the model without epistasis: 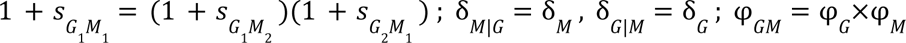

Initially, the genotype frequencies are:

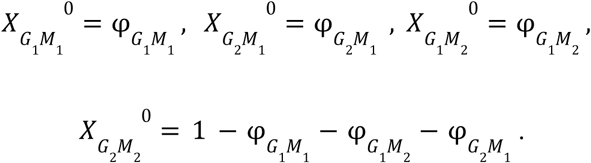

where φ*_G_* is the initial proportion of genotype *G*_1_, and is a model parameter.

The change in frequencies due to mutation is given by

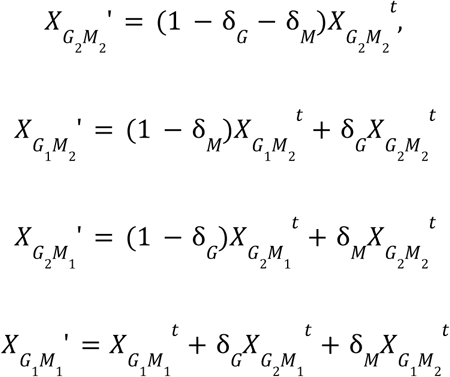

where *X*_*g*_ ’ is the frequency of genotype *g* after mutation.

The change in frequencies due to selection is given by

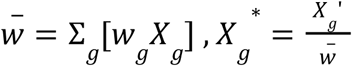

where *w*_*g*_ is the fitness of genotype *g*, and *w̄* is the population mean fitness. *X*_*g*_* represents the frequency of genotype *g* after selection.

The change in frequencies due to random genetic drift is given by

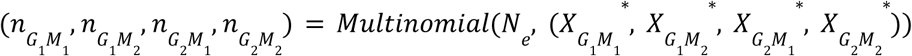

where *n*_*g*_ is the number of cells with genotype *g*.

The frequencies in the next generation are then given by

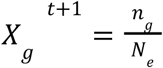

#### Predictive checks for the model with epistasis

To perform a predictive check, we inferred six model parameters using the model without epistasis for each of the replicates, calculated the value of the additional four parameters assuming independence of CNV reversions, simulated using these parameter estimates in the model with epistasis, and compared the simulation results to the empirical data. These predictions aligned well with the empirical data. Therefore, we concluded that the model without epistasis is sufficient and did not use the model with epistasis for parameter inference in the main text. In the future, the model with epistasis could be used to study results of experiments with epistasis between two-loci.

## Notes

### Competing Interest Statement

The authors have declared no competing interest.

### Summary of Updates

To complement our extensive short-read sequencing, we investigated multiple CNV revertant lineages using PacBio long-read sequencing (Figure 7). These data enabled us to resolve the precise structure of the partial CNV revertant and surprisingly revealed that it comprises a non-reciprocal translocation. Long read analysis also supports our finding that revertant strains are scarless, with segmental CNVs operating as ephemeral adaptive alleles leaving no trace of their transient existence. We propose a molecular mechanism for complete reversion (Figure 8). We investigated determinants of CNV strain fitness, and our results suggest that fitness cost is a joint function of the number of amplified genes and the individual effects of those genes (Figure 2C). We also undertook ploidy analysis of all CNV strains using DNA content staining, which revealed that four of the 15 CNV strains are diploid, whereas the others are haploid (Supplemental Figures S1, S2).

## References

1. Avecilla, G., Chuong, J.N., Li, F., Sherlock, G., Gresham, D. and Ram, Y., 2022. Neural networks enable efficient and accurate simulation-based inference of evolutionary parameters from adaptation dynamics. PLoS biology, 20(5), p.e3001633.

2. Avecilla, G., Spealman, P., Matthews, J., Caudal, E., Schacherer, J. and Gresham, D., 2023. Copy number variation alters local and global mutational tolerance. Genome Research, 33(8), pp.1340–1353.

3. Beach, R.R., Ricci-Tam, C., Brennan, C.M., Moomau, C.A., Hsu, P.H., Hua, B., Silberman, R.E., Springer, M. and Amon, A., 2017. Aneuploidy causes non-genetic individuality. Cell, 169(2), pp.229–242.

4. Ben-David, U. and Amon, A., 2020. Context is everything: aneuploidy in cancer. Nature Reviews Genetics, 21(1), pp.44–62.

5. Bonney, M.E., Moriya, H. and Amon, A., 2015. Aneuploid proliferation defects in yeast are not driven by copy number changes of a few dosage-sensitive genes. Genes & development, 29(9), pp.898–903.

6. Brown, C.J., Todd, K.M. and Rosenzweig, R.F., 1998. Multiple duplications of yeast hexose transport genes in response to selection in a glucose-limited environment. Molecular biology and evolution, 15(8), pp.931–942.

7. Campbell, D.A., Fogel, S. and Lusnak, K., 1975. Mitotic chromosome loss in a disomic haploid of Saccharomyces cerevisiae. Genetics, 79(3), pp.383–396.

8. Chang, S.L., Lai, H.Y., Tung, S.Y. and Leu, J.Y., 2013. Dynamic large-scale chromosomal rearrangements fuel rapid adaptation in yeast populations. PLoS genetics, 9(1), p.e1003232.

9. Chuong, J.N., Nun, N.B., Suresh, I., Matthews, J.C., De, T., Avecilla, G., Abdul-Rahman, F., Brandt, N., Ram, Y. and Gresham, D., 2025. Template switching during DNA replication is a prevalent source of adaptive gene amplification. eLife, 13, p.RP98934.

10. Clop, A., Vidal, O. and Amills, M., 2012. Copy number variation in the genomes of domestic animals. Animal genetics, 43(5), pp.503–517.

11. Connolly, J.J., Glessner, J.T., Almoguera, B., Crosslin, D.R., Jarvik, G.P., Sleiman, P.M. and Hakonarson, H., 2014. Copy number variation analysis in the context of electronic medical records and large-scale genomics consortium efforts. Frontiers in genetics, 5, p.51.

12. Cowell, A.N., Istvan, E.S., Lukens, A.K., Gomez-Lorenzo, M.G., Vanaerschot, M., Sakata-Kato, T., Flannery, E.L., Magistrado, P., Owen, E., Abraham, M. and LaMonte, G., 2018. Mapping the malaria parasite druggable genome by using in vitro evolution and chemogenomics. Science, 359(6372), pp.191–199.

13. Danecek, P., Bonfield, J.K., Liddle, J., Marshall, J., Ohan, V., Pollard, M.O., Whitwham, A., Keane, T., McCarthy, S.A., Davies, R.M. and Li, H., 2021. Twelve years of SAMtools and BCFtools. Gigascience, 10(2), p.giab008.

14. Dekel, E. and Alon, U., 2005. Optimality and evolutionary tuning of the expression level of a protein. Nature, 436(7050), pp.588–592.

15. Dhami, M.K., Hartwig, T. and Fukami, T., 2016. Genetic basis of priority effects: insights from nectar yeast. Proceedings of the Royal Society B: Biological Sciences, 283(1840), p.20161455.

16. Dolatabadian, A., Patel, D.A., Edwards, D. and Batley, J., 2017. Copy number variation and disease resistance in plants. Theoretical and Applied Genetics, 130(12), pp.2479–2490.

17. Dunham, M.J., Badrane, H., Ferea, T., Adams, J., Brown, P.O., Rosenzweig, F. and Botstein, D., 2002. Characteristic genome rearrangements in experimental evolution of Saccharomyces cerevisiae. Proceedings of the National Academy of Sciences, 99(25), pp.16144–16149.

18. Elde, N.C., Child, S.J., Eickbush, M.T., Kitzman, J.O., Rogers, K.S., Shendure, J., Geballe, A.P. and Malik, H.S., 2012. Poxviruses deploy genomic accordions to adapt rapidly against host antiviral defenses. Cell, 150(4), pp.831–841.

19. Freeling, M., Scanlon, M.J. and Fowler, J.E., 2015. Fractionation and subfunctionalization following genome duplications: mechanisms that drive gene content and their consequences. Current Opinion in Genetics & Development, 35, pp.110–118.

20. Galanti, L., Shasha, D. and Gunsalus, K.C., 2021. Pheniqs 2.0: accurate, high-performance Bayesian decoding and confidence estimation for combinatorial barcode indexing. BMC bioinformatics, 22(1), p.359.

21. “GENEFLOW.” 2023. New York University Center for Genomics and System Biology Genomics Core, GitHub Repository. https://github.com/gencorefacility/GENEFLOW; New York University.

22. Gerstein, A.C. and Berman, J., 2015. Shift and adapt: the costs and benefits of karyotype variations. Current opinion in Microbiology, 26, pp.130–136.

23. Golzio, C. and Katsanis, N., 2013. Genetic architecture of reciprocal CNVs. Current opinion in genetics & development, 23(3), pp.240–248.

24. Good, B.H., McDonald, M.J., Barrick, J.E., Lenski, R.E. and Desai, M.M., 2017. The dynamics of molecular evolution over 60,000 generations. Nature, 551(7678), pp.45–50.

25. Greenblum, S., Carr, R. and Borenstein, E., 2015. Extensive strain-level copy-number variation across human gut microbiome species. Cell, 160(4), pp.583–594.

26. Gresham, D., Desai, M.M., Tucker, C.M., Jenq, H.T., Pai, D.A., Ward, A., DeSevo, C.G., Botstein, D. and Dunham, M.J., 2008. The repertoire and dynamics of evolutionary adaptations to controlled nutrient-limited environments in yeast. PLoS genetics, 4(12), p.e1000303.

27. Gresham, D., Usaite, R., Germann, S.M., Lisby, M., Botstein, D. and Regenberg, B., 2010. Adaptation to diverse nitrogen-limited environments by deletion or extrachromosomal element formation of the GAP1 locus. Proceedings of the National Academy of Sciences, 107(43), pp.18551–18556.

28. Gresham, D. and Dunham, M.J., 2014. The enduring utility of continuous culturing in experimental evolution. Genomics, 104(6), pp.399–405.

29. Gresham, D. and Hong, J., 2015. The functional basis of adaptive evolution in chemostats. FEMS microbiology reviews, 39(1), pp.2–16.

30. Hoffman, C.S. and Winston, F., 1987. A ten-minute DNA preparation from yeast efficiently releases autonomous plasmids for transformation of Escherichia coli. Gene, 57(2-3), pp.267–272.

31. Hong, J. and Gresham, D., 2014. Molecular specificity, convergence and constraint shape adaptive evolution in nutrient-poor environments. PLoS genetics, 10(1), p.e1004041.

32. Hong, J., Brandt, N., Abdul-Rahman, F., Yang, A., Hughes, T. and Gresham, D., 2018. An incoherent feedforward loop facilitates adaptive tuning of gene expression. Elife, 7, p.e32323.

33. Horiuchi, T., Horiuchi, S. and Novick, A., 1963. The genetic basis of hyper-synthesis of β-galactosidase. Genetics, 48(2), p.157.

34. Iantorno, S.A., Durrant, C., Khan, A., Sanders, M.J., Beverley, S.M., Warren, W.C., Berriman, M., Sacks, D.L., Cotton, J.A. and Grigg, M.E., 2017. Gene expression in Leishmania is regulated predominantly by gene dosage. MBio, 8(5), pp.10–1128.

35. Innan, H. and Kondrashov, F., 2010. The evolution of gene duplications: classifying and distinguishing between models. Nature Reviews Genetics, 11(2), pp.97–108.

36. Iskow, R.C., Gokcumen, O., Abyzov, A., Malukiewicz, J., Zhu, Q., Sukumar, A.T., Pai, A.A., Mills, R.E., Habegger, L., Cusanovich, D.A. and Rubel, M.A., 2012. Regulatory element copy number differences shape primate expression profiles. Proceedings of the National Academy of Sciences, 109(31), pp.12656–12661.

37. Kao, K.C. and Sherlock, G., 2008. Molecular characterization of clonal interference during adaptive evolution in asexual populations of Saccharomyces cerevisiae. Nature genetics, 40(12), pp.1499–1504.

38. Kohanovski, I., Pontz, M., Vande Zande, P., Selmecki, A., Dahan, O., Pilpel, Y., Yona, A.H. and Ram, Y., 2024. Aneuploidy can be an evolutionary diversion on the path to adaptation. Molecular biology and evolution, 41(3), p.msae052.

39. Kondrashov, F.A., 2012. Gene duplication as a mechanism of genomic adaptation to a changing environment. Proceedings of the Royal Society B: Biological Sciences, 279(1749), pp.5048–5057.

40. Koszul, R., Dujon, B. and Fischer, G., 2006. Stability of large segmental duplications in the yeast genome. Genetics, 172(4), pp.2211–2222.

41. Lauer, S., Avecilla, G., Spealman, P., Sethia, G., Brandt, N., Levy, S.F. and Gresham, D., 2018. Single-cell copy number variant detection reveals the dynamics and diversity of adaptation. PLoS Biology, 16(12), p.e3000069.

42. Lenski, R.E., Rose, M.R., Simpson, S.C. and Tadler, S.C., 1991. Long-term experimental evolution in Escherichia coli. I. Adaptation and divergence during 2,000 generations. The American Naturalist, 138(6), pp.1315–1341.

43. Li, H., 2016. Minimap and miniasm: fast mapping and de novo assembly for noisy long sequences. Bioinformatics, 32(14), pp.2103–2110.

44. Li, H., 2018. Minimap2: pairwise alignment for nucleotide sequences. Bioinformatics, 34(18), pp.3094–3100.

45. Linder, R.A., Greco, J.P., Seidl, F., Matsui, T. and Ehrenreich, I.M., 2017. The stress-inducible peroxidase TSA2 underlies a conditionally beneficial chromosomal duplication in Saccharomyces cerevisiae. G3: Genes, Genomes, Genetics, 7(9), pp.3177–3184.

46. Lukow, D.A., Sausville, E.L., Suri, P., Chunduri, N.K., Wieland, A., Leu, J., Smith, J.C., Girish, V., Kumar, A.A., Kendall, J. and Wang, Z., 2021. Chromosomal instability accelerates the evolution of resistance to anti-cancer therapies. Developmental cell, 56(17), pp.2427–2439.

47. Makanae, K., Kintaka, R., Makino, T., Kitano, H. and Moriya, H., 2013. Identification of dosage-sensitive genes in Saccharomyces cerevisiae using the genetic tug-of-war method. Genome research, 23(2), pp.300–311.

48. Myhre, S., Lingjærde, O.C., Hennessy, B.T., Aure, M.R., Carey, M.S., Alsner, J., Tramm, T., Overgaard, J., Mills, G.B., Børresen-Dale, A.L. and Sørlie, T., 2013. Influence of DNA copy number and mRNA levels on the expression of breast cancer related proteins. Molecular oncology, 7(3), pp.704–718.

49. Ohno, S., 2013. Evolution by gene duplication. Springer Science & Business Media.

50. Payen, C., Sunshine, A.B., Ong, G.T., Pogachar, J.L., Zhao, W. and Dunham, M.J., 2016. High-throughput identification of adaptive mutations in experimentally evolved yeast populations. PLoS genetics, 12(10), p.e1006339.

51. “Picard Toolkit.” 2019. Broad Institute, GitHub Repository. https://broadinstitute.github.io/picard/; Broad Institute.

52. Quinlan, A.R. and Hall, I.M., 2010. BEDTools: a flexible suite of utilities for comparing genomic features. Bioinformatics, 26(6), pp.841–842.

53. Reams, A.B., Kofoid, E., Savageau, M. and Roth, J.R., 2010. Duplication frequency in a population of Salmonella enterica rapidly approaches steady state with or without recombination. Genetics, 184(4), pp.1077–1094.

54. Rice, A.M. and McLysaght, A., 2017. Dosage-sensitive genes in evolution and disease. BMC biology, 15(1), p.78.

55. Robinson, J.T., Thorvaldsdóttir, H., Winckler, W., Guttman, M., Lander, E.S., Getz, G. and Mesirov, J.P., 2011. Integrative genomics viewer. Nature biotechnology, 29(1), pp.24–26.

56. Robinson, D., Place, M., Hose, J., Jochem, A. and Gasch, A.P., 2021. Natural variation in the consequences of gene overexpression and its implications for evolutionary trajectories. Elife, 10, p.e70564.

57. Robinson, D., Vanacloig-Pedros, E., Cai, R., Place, M., Hose, J. and Gasch, A.P., 2023. Gene-by-environment interactions influence the fitness cost of gene copy-number variation in yeast. G3: Genes, Genomes, Genetics, 13(10), p.jkad159.

58. Rojas, J., Hose, J., Dutcher, H.A., Place, M., Wolters, J.F., Hittinger, C.T. and Gasch, A.P., 2024. Comparative modeling reveals the molecular determinants of aneuploidy fitness cost in a wild yeast model. Cell Genomics, 4(10).

59. Rutledge, S.D., Douglas, T.A., Nicholson, J.M., Vila-Casadesús, M., Kantzler, C.L., Wangsa, D., Barroso-Vilares, M., Kale, S.D., Logarinho, E. and Cimini, D., 2016. Selective advantage of trisomic human cells cultured in non-standard conditions. Scientific reports, 6(1), p.22828.

60. Sheltzer, J.M., Torres, E.M., Dunham, M.J. and Amon, A., 2012. Transcriptional consequences of aneuploidy. Proceedings of the National Academy of Sciences, 109(31), pp.12644–12649.

61. Smith, J.M. and Haigh, J., 1974. The hitch-hiking effect of a favourable gene. Genetics Research, 23(1), pp.23–35.

62. Sonti, R.V. and Roth, J.R., 1989. Role of gene duplications in the adaptation of Salmonella typhimurium to growth on limiting carbon sources. Genetics, 123(1), pp.19–28.

63. Spealman, P., Burrell, J. and Gresham, D., 2020. Inverted duplicate DNA sequences increase translocation rates through sequencing nanopores resulting in reduced base calling accuracy. Nucleic Acids Research, 48(9), pp.4940–4945.

64. Spealman, P., De, T., Chuong, J.N. and Gresham, D., 2023. Best practices in microbial experimental evolution: using reporters and long-read sequencing to identify copy number variation in experimental evolution. Journal of Molecular Evolution, 91(3), pp.356–368.

65. Spealman, P., de Santana, C., De, T. and Gresham, D., 2025. Multilevel gene expression changes in lineages containing adaptive copy number variants. Molecular Biology and Evolution, 42(2), p.msaf005.

66. Sunshine, A.B., Payen, C., Ong, G.T., Liachko, I., Tan, K.M. and Dunham, M.J., 2015. The fitness consequences of aneuploidy are driven by condition-dependent gene effects. PLoS biology, 13(5), p.e1002155.

67. Tang, Y.C. and Amon, A., 2013. Gene copy-number alterations: a cost-benefit analysis. Cell, 152(3), pp.394–405.

68. Todd, R.T., Braverman, A.L. and Selmecki, A., 2018. Flow cytometry analysis of fungal ploidy. Current protocols in microbiology, 50(1), p.e58.

69. Todd, R.T. and Selmecki, A., 2020. Expandable and reversible copy number amplification drives rapid adaptation to antifungal drugs. Elife, 9, p.e58349.

70. Tomanek, I., Grah, R., Lagator, M., Andersson, A.M.C., Bollback, J.P., Tkačik, G. and Guet, C.C., 2020. Gene amplification as a form of population-level gene expression regulation. Nature ecology & evolution, 4(4), pp.612–625.

71. Tomanek, I and Guet, C.C., 2022. Adaptation dynamics between copy-number and point mutations. eLife, 11.

72. Torres, E.M., Sokolsky, T., Tucker, C.M., Chan, L.Y., Boselli, M., Dunham, M.J. and Amon, A., 2007. Effects of aneuploidy on cellular physiology and cell division in haploid yeast. science, 317(5840), pp.916–924.

73. Tsai, H.J., Nelliat, A.R., Choudhury, M.I., Kucharavy, A., Bradford, W.D., Cook, M.E., Kim, J., Mair, D.B., Sun, S.X., Schatz, M.C. and Li, R., 2019. Hypo-osmotic-like stress underlies general cellular defects of aneuploidy. Nature, 570(7759), pp.117–121.

74. Tung, S., Bakerlee, C.W., Phillips, A.M., Nguyen Ba, A.N. and Desai, M.M., 2021. The genetic basis of differential autodiploidization in evolving yeast populations. G3, 11(8), p.jkab192.

75. Veitia, R.A., Bottani, S. and Birchler, J.A., 2008. Cellular reactions to gene dosage imbalance: genomic, transcriptomic and proteomic effects. Trends in Genetics, 24(8), pp.390–397.

76. Wagner, A., 2005. Energy constraints on the evolution of gene expression. Molecular biology and evolution, 22(6), pp.1365–1374.

77. Warringer, J., Zörgö, E., Cubillos, F.A., Zia, A., Gjuvsland, A., Simpson, J.T., Forsmark, A., Durbin, R., Omholt, S.W., Louis, E.J. and Liti, G., 2011. Trait variation in yeast is defined by population history. PLoS genetics, 7(6), p.e1002111.

78. Yang, F., Todd, R.T., Selmecki, A., Jiang, Y.Y., Cao, Y.B. and Berman, J., 2021. The fitness costs and benefits of trisomy of each Candida albicans chromosome. Genetics, 218(2), p.iyab056.

79. Yona, A.H., Manor, Y.S., Herbst, R.H., Romano, G.H., Mitchell, A., Kupiec, M., Pilpel, Y. and Dahan, O., 2012. Chromosomal duplication is a transient evolutionary solution to stress. Proceedings of the National Academy of Sciences, 109(51), pp.21010–21015.

80. Zarrei, M., MacDonald, J.R., Merico, D. and Scherer, S.W., 2015. A copy number variation map of the human genome. Nature reviews genetics, 16(3), pp.172–183.

81. Zhang, F., Gu, W., Hurles, M.E. and Lupski, J.R., 2009. Copy number variation in human health, disease, and evolution. Annual review of genomics and human genetics, 10(1), pp.451–481.

82. Zhong, S., Khodursky, A., Dykhuizen, D.E. and Dean, A.M., 2004. Evolutionary genomics of ecological specialization. Proceedings of the National Academy of Sciences, 101(32), pp.11719–11724.

83. Żmieńko, A., Samelak, A., Kozłowski, P. and Figlerowicz, M., 2014. Copy number polymorphism in plant genomes. Theoretical and applied genetics, 127(1), pp.1–18.

